# Chemokine Signatures Of Pathogen-Specific T Cells I: Effector T Cells

**DOI:** 10.1101/2020.02.05.935502

**Authors:** Jens Eberlein, Bennett Davenport, Tom T. Nguyen, Francisco Victorino, Kevin Jhun, Verena van der Heide, Maxim V. Kuleshov, Avi Ma’ayan, Ross Kedl, Dirk Homann

**Affiliations:** Barbara Davis Center for Childhood Diabetes, University of Colorado Denver, Aurora CO; Integrated Department of Immunology, University of Colorado Denver and National Jewish Health, Denver CO; Department of Anesthesiology, University of Colorado Denver, Aurora, CO; Diabetes Obesity Metabolism Institute, Icahn School of Medicine at Mount Sinai, New York, NY; Immunology Institute, Icahn School of Medicine at Mount Sinai, New York, NY; Department of Pharmacological Sciences, Icahn School of Medicine at Mount Sinai, New York, NY; Mount Sinai Center for Bioinformatics, Icahn School of Medicine at Mount Sinai, New York, NY

## Abstract

The choreography of complex immune responses, including the priming, differentiation, and modulation of specific effector T cell populations generated in the immediate wake of an acute pathogen challenge, is in part controlled by chemokines, a large family of mostly secreted molecules involved in chemotaxis and other patho/physiological processes. T cells are both responsive to varied chemokine cues and a relevant source for certain chemokines themselves. Yet the actual range, regulation, and role of effector T cell-derived chemokines remains incompletely understood. Here, using different *in vivo* models of viral and bacterial infection as well as protective vaccination, we have defined the entire spectrum of chemokines produced by pathogen-specific CD8^+^ and CD4^+^T effector cells, and delineated several unique properties pertaining to the temporospatial organization of chemokine expression patterns, synthesis and secretion kinetics, and cooperative regulation. Collectively, our results position the “T cell chemokine response” as a notably prominent, largely invariant yet distinctive force at the forefront of pathogen-specific effector T cell activities, and establish novel practical and conceptual approaches that may serve as a foundation for future investigations into role of T cell-produced chemokines in infectious and other diseases.

## INTRODUCTION

The immune system is a distributed network of organs, tissues, cells and extracellular factors. Functional integration of these components faces a particular challenge since the principal sentinels, regulators and effectors of immune function are often highly mobile single cells. The controlled spatiotemporal positioning of these cells is achieved by adhesion molecules such as integrins and selectins as well as chemokines and their receptors that function as a “molecular address” system in the coordination of cellular traffic in specific tissue microenvironments [1–4]. The defining function of chemokines (*chemo*attractant cyto*kines*), demonstrated in numerous *in vitro* experiments, is their capacity to induce the directed migration of locomotive cells by establishing a spatial gradient. However, chemokines exhibit a host of additional functions including control of lymphopoiesis and lymphoid organogenesis, alterations of leukocyte adhesive properties by modulation of integrins as well as regulation of lymphocyte differentiation, proliferation, survival, cytokine release and degranulation [1, 3, 5–7]. Given this functional diversity, chemokines have been implicated in a wide variety of pathological states such as infectious disease and cancer, autoimmunity, allergy and transplant rejection [7–12].

The family of chemokines comprises a large number of mainly secreted molecules that share a defining tetracysteine motif and can be classified according to structural criteria, functional properties (“homeostatic” *vs.* “inflammatory”) and genomic organization [13–15]. Among the many different cell types capable of chemokine production, pathogen-specific T cells were identified as a relevant source over two decades ago [16]. However, while the T cell-produced chemokines CCL3/4/5 have received considerable attention as competitive inhibitors of HIV binding to its co-receptor CCR5 [17–19], an inclusive perspective on specific T cell-produced chemokines has not been established, a likely consequence of both an experimental and conceptual emphasis on chemokine action *on* T cells rather than chemokine production *by* T cells [20–23].

In the more circumscribed context of pathogen-specific effector T cell (T_E_) immunity, *i.e.* T cell responses generated in the immediate wake of an acute pathogen challenge and the topic of the present investigations, murine models of infectious disease have by and large confirmed the prodigious CCL3/4/5 production capacity of T_E_ populations. For example, Dorner *et al.* demonstrated that CCL3/4/5 as well as XCL1 are readily synthesized by CD8^+^ but not CD4^+^T_E_ specific for the bacterium *L. monocytogenes* (LM), are co-expressed with IFNγ, and thus may constitute a family of “type 1 cytokines” [24]. Moreover, CCL3-deficient but not wild-type (wt) LM-specific CD8^+^T_E_, after transfer into naïve wt recipients, failed to protect against a lethal LM infection, to this date one of the most striking phenotypes reported for a T cell-specific chemokine deficiency [25]. Abundant CCL3/4/5 is also made by CD8^+^T_E_, and to a lesser extent by CD4^+^T_E_, generated in response to acute infection with lymphocytic choriomeningitis virus (LCMV) [26]. In the related LCMV model of lethal choriomeningitis, CCL3/4/5 secretion by CD8^+^T_E_ has been associated with the recruitment of pathogenic myelomonocytic cells into the CNS and lethal choriomeningitis [27] but the precise role of these chemokines remains to be determined given that mice deficient for CCL3 or CCR5 (only receptor for CCL4 that also binds CCL3/5) are not protected from fatal disease [28]. Even during the initial stages of T cell priming, CCL3/4 production by activated CD4^+^ or CD8^+^ T cells (induced by peptide immunization or vaccinia virus infection, respectively) contributes to the effective spatiotemporal organization of T and dendritic cell interactions [29, 30]. A similar role has most recently also been demonstrated for CD8^+^T cell-derived XCL1 [30] and, following an earlier report that CD8^+^T cell-secreted XCL1 is required for optimal proliferative expansion of allogeneic CD8^+^T_E_ [31], mice lacking XCR1 (the sole XCL1 receptor) were shown to generate reduced LM-specific CD8^+^T_E_ responses associated with delayed bacterial control [32]. Collectively, these observations demonstrate that pathogen-specific CD8^+^ and CD4^+^T cells, beyond their responsiveness to numerous varied chemokine cues, are themselves a relevant source for select chemokines that exert non-redundant effects on the development of effective T_E_ responses and, in some cases, efficient pathogen control.

The complete range of chemokines produced by pathogen-specific CD8^+^ and CD4^+^T_E_, however, has not yet been defined, and the respective expression patterns of T cell-derived chemokines, their co-regulation as well as synthesis and secretion kinetics remain incompletely understood. Here, we have addressed these issues in a series of complementary investigations that chiefly rely on the use of stringently characterized chemokine-specific antibodies that permit the flow cytometry-(FC-) based detection of practically all (37 out of 38) murine chemokines at the single-cell level [33]. Our results demonstrate that production of chemokines by pathogen-specific CD8^+^ and CD4^+^T_E_ constitutes a restricted (CCL1, CCL3, CCL4, CCL5, CCL9/10 and XCL1), remarkably prominent, uniquely regulated, integral and consistent component of the T_E_ response across different infectious disease models and protective vaccination; together, these properties position mature T_E_-derived chemokines at the forefront of coordinated host pathogen defenses.

## RESULTS

### Broad survey of T cell-produced chemokines

To delineate the principal spectrum of chemokines synthesized by activated T cells, spleen cells obtained from unmanipulated wt mice were stimulated for 5h with PMA/ionomycin and interrogated by flow cytometry (FC) for production of IFNγ and 37 individual chemokines using a stringently validated panel of chemokine-specific antibodies [33]. Robust induction was observed for CCL3, CCL4 and XCL1 and to a lesser extent also CCL5; 1-2% of T cells synthesized CXCL2; and very small subsets produced CCL1 or CCL9/10 (***Fig.S1A-C***). CCL3, CCL4, CCL5 and XCL1 production was particularly prominent in the IFNγ^+^ T cell subset (40-80% co-expression) and despite their low frequency, CCL1- and CCL9/10-expressing T cells were also enriched in the IFNγ^+^ population (3-4% co-expression); in contrast, no such enrichment was observed for CXCL2 (***Fig.S1A/C***). Although T cells have previously been described as a source for these chemokines in various experimental scenarios, we note that the comprehensive nature of our screen, within the limits of specific experimental constraints and the sensitivity afforded by chemokine FC [33], can apparently rule out 30 other chemokines as potential products of highly activated T cells.

### Defining the complete spectrum of chemokines produced by virus-specific CD8^+^T_E_

In order to refine these analyses within the context of infectious diseases and to define the complete range of chemokines produced by pathogen-specific CD8^+^T^E^, we first employed the established “p14 chimera” system to quantify chemokine mRNA and protein expression by virus-specific CD8^+^T^E^ with a combination of gene arrays and FC-based assays [33, 34]. In brief, p14 chimeras were generated by transducing congenic B6 mice with a trace population of naïve TCR transgenic CD8^+^T cells (p14 T^N^) specific for the dominant LCMV-GP^33-41^ determinant; after challenge with LCMV, p14 T^E^ populations rapidly differentiate, expand and contribute to efficient virus control before contracting and developing into p14 memory T cells (T^M^) ∼6 weeks later [34–36]. At the peak of the effector phase (d8), p14 T^E^ were purified, RNA was extracted either immediately or after a 3h *in vitro* TCR stimulation, and processed for gene array hybridization. Overall, 10 chemokine mRNA species were detectable in p14 T^E^ evaluated *ex vivo* and*/*or after TCR stimulation, and their expression patterns could be allocated to three groups (***Fig.1A***): 1., absence of *ex vivo* detectable mRNA but robust transcription after TCR stimulation (*Ccl1* and *Xcl1*); 2., constitutive mRNA expression that significantly increased upon TCR engagement (*Ccl3*, *Ccl4*, *Ccl9/10* and *Cxcl10*); and 3., chemokine mRNA species that were slightly downregulated by TCR activation (*Ccl5*, *Ccl6*, *Ccl25*, and *Ccl27*); a list of all murine chemokine genes and gene array IDs is found in ***Fig.S2A***. We also quantified chemokine mRNA expression for the known members of the related chemokine-like factor superfamily (CKLFSF) [37] (***Fig.S2B***). Four out of 10 *Cklfsf* mRNA species were detected in p14 T^E^ but none were in- or decreased upon TCR stimulation. Information about the biological function of CKLFSF members remains limited and is centered around the pleiotropic effects of CKLF1 which may be produced by human T cells after prolonged *in vitro* stimulation [38, 39]. At the present stage, we have refrained from a further analysis of this gene family.

**Figure 1.**
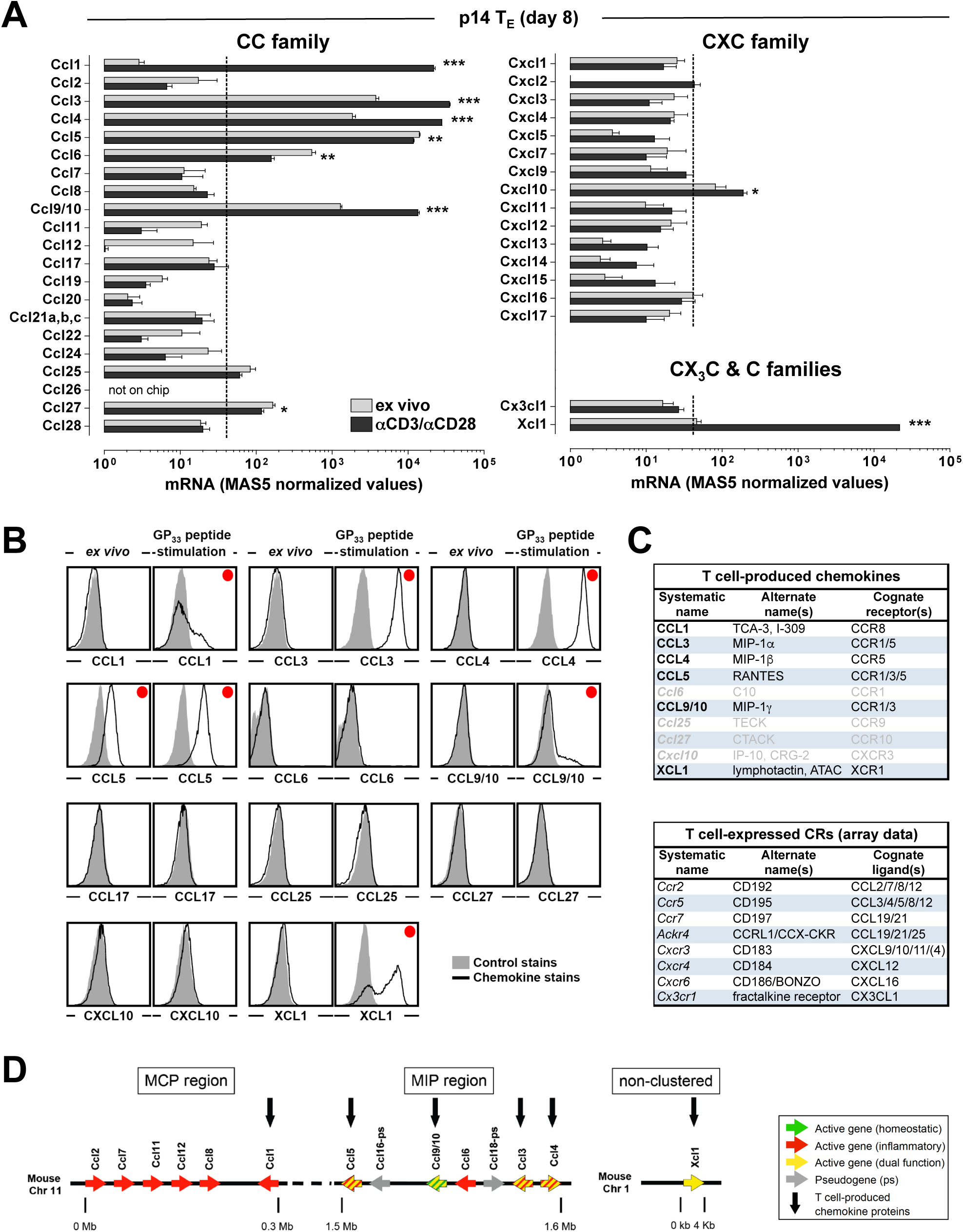
Chemokine mRNA and protein expression by virus-specific CD8^+^T_E_. **A.,** p14 T_E_ (day 8) were obtained from spleens of LCMV-infected p14 chimeras, enriched to >99% purity and processed for RNA extraction (either immediately or after 3h αCD3/αCD28 stimulation) and gene array analysis (n=3 individual mice). The bar diagrams display MAS5-normalized values of chemokine mRNA expression of p14 T_E_ analyzed *ex vivo* (gray bars) or after TCR stimulation (black bars); statistically significant differences are indicated by asterisks. The broken line indicates the detection threshold set at a MAS5 value of 40; coverage: 39/40 chemokines (*Ccl26* not on chip). **B.,** p14 T_E_ (d8) were analyzed for chemokine protein expression *ex vivo* or after 5h stimulation with GP_33_ peptide. Histograms are gated on p14 cells (gray histograms: control stains, black tracings: indicated chemokine stains; red dots identify panels demonstrating detectable chemokine expression). **C.,** summary of p14 T_E_- and T_M_-expressed chemokines and chemokine receptors; gray font indicates presence of mRNA in the absence of constitutive or induced protein expression. **D.,** genomic organization of murine chemokine genes transcribed and translated by T cells (modified after ref. [14]). The genes for 4/6 chemokine produced by T cells (*Ccl3/4/5* and *Ccl9/10*) are found in the MIP region on mouse chromosome 11; the *Ccl1* gene is located in the MCP region but rather distantly related to other members of the MCP group, and the non-clustered *Xcl1* gene is found on chromosome 1. Arrows indicate chemokine genes and their transcriptional orientation; colors identify homeostatic (green), inflammatory (red) and dual function (yellow) chemokine genes; gray arrows indicate pseudogenes. Based on our results reported here and in ref. [33] we propose to classify the CCL3/4/5 and 9/10 as “dual function” chemokines rather than simply “inflammatory” (CCL3/4/5) or “homeostatic” (CCL9/10).

Traditional gene array analyses are fraught with several limitations including measurement of mRNA levels across entire, albeit purified, cell populations rather than individual cells; difficulties in directly comparing mRNA expression between different mRNA species; and the impossibility to predict if mRNA is in fact translated. We therefore deployed chemokine FC to the interrogation of p14 T^E^ and found that specific TCR stimulation with GP^33^ peptide induced CCL3, CCL4 and CCL5 expression in practically all p14 T^E^ while CCL1, CCL9/10 and XCL1 production was restricted to p14 T^E^ subsets; neither *Ccl6*, *Ccl25*, *Ccl27*, *Cxcl10* nor *Cxcl2* or any other chemokine RNA was translated. Of note, CCL5 was also detectable directly *ex vivo* and is thus the only chemokine expressed in the absence of TCR activation (***Fig.1B***). The robust induction of CCL1 and CCL9/10 in p14 T^E^ contrasts with their more limited expression in our initial T cell survey (***Fig.S1A-C***). To resolve these discrepancies, we quantified CCL1 and CCL9/10 production by CD8^+^T cells from LCMV-immune mice in response to stimulation with peptide, αCD3/αCD28 or PMA/ionomycin. Interestingly, >20% of IFNγ^+^ CD8^+^T cells also synthesized CCL1 or CCL9/10 after activation with peptide or αCD3/αCD28 but fewer than 5% produced these chemokines in response to PMA/ionomycin stimulation (***Fig.S1D***). The reason for the only sparse CCL1 and CCL9/10 induction after PMA/ionomycin treatment remains unclear but emphasizes important limitations associated with this widely used T cell stimulation protocol. In summary, six mRNA species found *ex vivo* and/or after brief TCR engagement in virus-specific p14 T^E^ serve as templates for induced protein synthesis and, in the case of *Ccl5*, also for effective constitutive translation (*Fig.1B/C*). The genes of four of these six chemokines (*Ccl3/4/5* and *9/10*) are clustered in the “MIP region” on murine chromosome 11 with an additional gene (*Ccl1*) immediately adjacent in the “MCP region”; the *Xcl1* gene is unclustered and located on chromosome 1 (***Fig.1D***).

Lastly, transcriptional profiling of p14 T^E^-expressed chemokine receptors revealed a prominent presence of *Ccr2*, *Cxcr3*, *Cxcr4*, *Cx3cr1* and *Ccrl1* (all of which were significantly downregulated upon activation); *Ccr5* and *Cxcr6* (slightly increased after stimulation); and low levels of *Ccr7* that remained unaffected by TCR engagement (***Fig.1C*** and not shown). It therefore appears that CCR5 is the only receptor that may sensitize p14 T^E^ to potential auto- or paracrine actions of T cell-produced chemokines themselves (i.e., CCL3/4/5).

### Constitutive and induced chemokine expression profiles of endogenously generated LCMV-specific CD8*^+^* and CD4^+^T^E^

Extending our findings from the p14 chimera system to endogenously generated T^E^, direct *ex vivo* analyses of the dominant D^b^NP^396+^ CD8^+^T^E^ population in LCMV-infected B6 mice demonstrated patterns comparable to p14 T^E^ in that constitutive chemokine expression was largely limited to CCL5. The small subsets of specific CD8^+^T^E^ showing weak CCL1/3/4 (but no CCL9/10 or XCL1) staining suggest that their expression, in contrast to IFNγ and other cytokines [40], can be maintained somewhat longer after cessation of TCR activation (***Fig.2A***). Constitutive CCL5 expression, on the other hand, is a general feature of LCMV-specific CD8^+^T^E^ as based on analyses of additional epitope-specific CD8^+^T^E^ and the inclusion of LCMV-infected *Ccl5*-deficient mice as a negative control (we also observed a slight reduction of *ex vivo* detectable CCL5 in *Ccl5* heterozygous mice indicative of a modest gene dosage effect) (***Fig.2A***), and appears to be in fact unique for cytokines at large, since no other effector molecules readily detected in re-stimulated CD8^+^T^E^ (IFNγ, TNFα, IL-2, GM-CSF, CD40L) are expressed in a constitutive fashion (***Fig.S3A/B***). Rather, CCL5 expression resembles that of constituents of the granzyme/perforin pathway [41–44] (***Fig.S3A/C***). Similar to CD8^+^T^E^, LCMV-specific CD4^+^T^E^ cells also contained *ex vivo* detectable CCL5 albeit only in a subset (∼60%) and at lower levels (***Fig.2B***).

**Figure 2.**
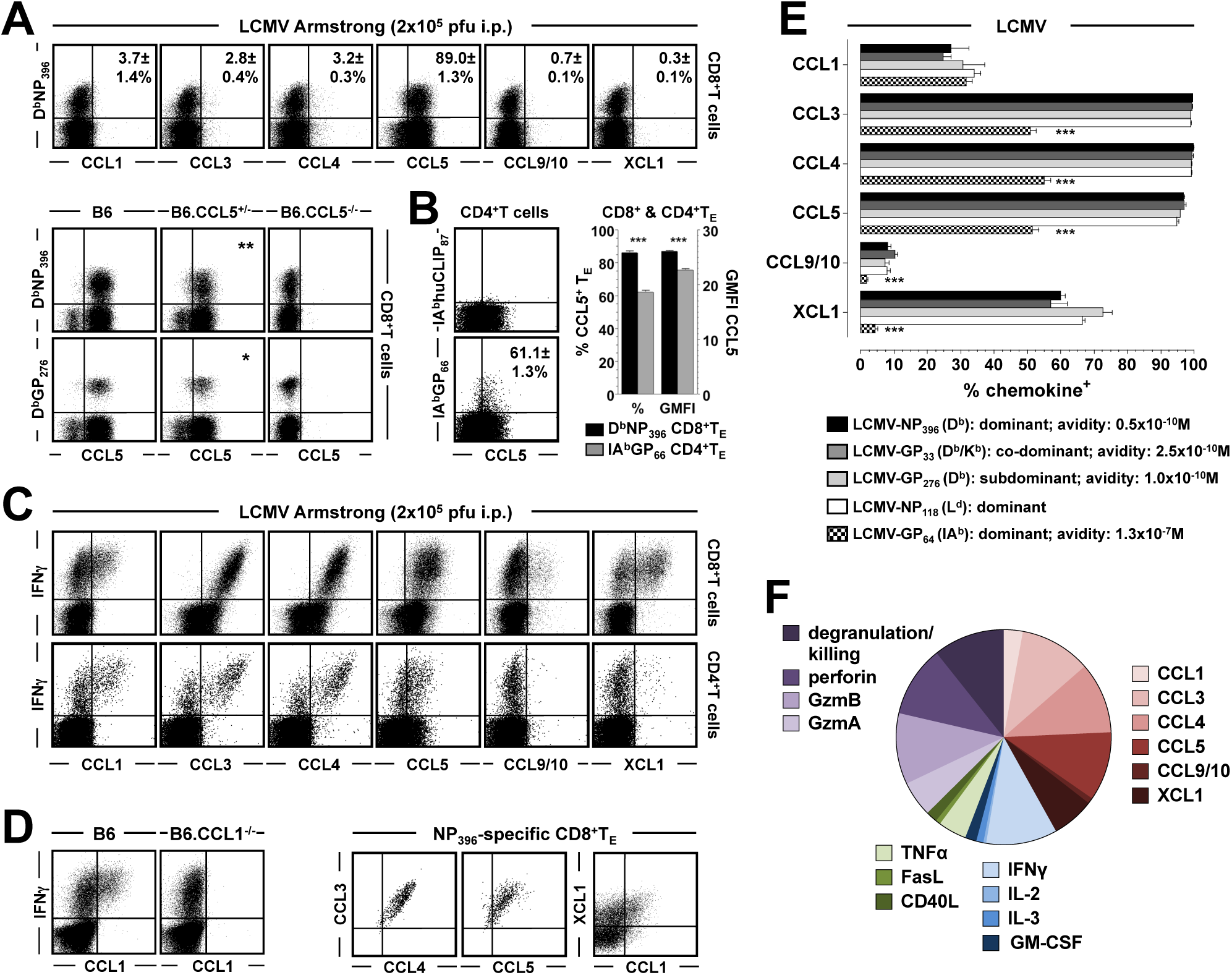
Constitutive and induced chemokine expression by LCMV-specific CD8^+^ and CD4^+^T_E_. **A.,** top row: Endogenously generated LCMV-specific CD8^+^T_E_ (d8) were analyzed directly *ex vivo* by chemokine FC (plots gated on splenic CD8^+^T cells); values indicate SEM of chemokine^+^ subsets among NP_396_-specific CD8^+^T_E_. Middle and bottom rows: constitutive CCL5 expression by LCMV-specific CD8^+^T_E_ subsets generated by LCMV-infected B6, B6.CCL5^+/-^ and B6.CCL5^-/-^ mice. **B.,** *ex vivo* detectable CCL5 expression by LCMV-specific CD4^+^T_E_. The adjacent bar diagram compares the fractions (%) and CCL5 expression levels (GMFI: geometric mean of fluorescence intensity) of CCL5^+^ specific CD8^+^ and CD4^+^T cells; statistical differences are indicated by asterisks. For the purpose of this direct comparison, MHC-I and -II tetramer stains were performed under the same experimental conditions (90min incubation at 37°C). **C.,** induced chemokine production by NP_396_-specific CD8^+^ (top row) and GP_64_-specific CD4^+^ (bottom row) T cells as determined after 5h *in vitro* peptide stimulation culture. **D.,** top: induced CCL1 expression following NP_396_ peptide stimulation of d8 spleen cells from LCMV-infected B6 and B6.CCL1^-/-^ mice. Bottom: chemokine co-expression by NP_396_-specific CD8^+^T_E_ (plots gated on IFNγ^+^ CD8^+^T_E_). **E.,** summary of induced chemokine expression by LCMV-specific T_E_ subsets stratified according epitope specificity; their restriction elements, relative size (immunodominance) and functional avidities (peptide concentration required to induce IFNγ production in 50% of a given epitope-specific population) are indicated. Significant differences between chemokine-expressing CD8^+^ and CD4^+^T_E_ subsets are indicated by asterisks. **F.,** the composition of the NP_396_-specific CD8^+^T_E_ response (d8) was assessed by quantification of subsets expressing individual constitutive (GzmA/B and perforin) or inducible (all other including CCL5) effector activities, and the pie chart depicts the sum and relative distribution thereof. All data (SEM) are representative for multiple experiments comprising groups of 3-5 individual mice.

Upon *in vitro* re-stimulation, both CD8^+^ and CD4^+^T^E^ rapidly synthesized the same six chemokines induced in p14 T^E^ but no other chemokines (***Fig.2C*** and not shown). Accordingly, CCL1, CCL9/10 and XCL1 production by specific CD8^+^T^E^ was mostly restricted to a subset of IFNγ^+^ cells whereas CCL3/4/5 were produced by virtually all epitope-specific CD8^+^T^E^. The patterns of induced chemokine synthesis further indicated the existence of particularly potent CD8^+^T^E^ populations as demonstrated by the co-expression of high CCL3/4/5 levels and the relative restriction of CCL1 production to a subset of XCL1^+^ CD8^+^T^E^ (***Fig.2D***). While the chemokine profiles of specific CD4^+^T^E^ were qualitatively similar, induced CCL3/4/5 production was confined to a subset (∼60%) and very few cells produced CCL9/10 or XCL1 (***Fig.2C***). ***Fig.2E*** summarizes these findings by displaying the fraction of chemokine^+^ LCMV-specific T^E^ stratified according to the MHC restriction element. Since the different epitope-specific T^E^ populations not only differed according to immunodominance but also activation threshold (***Fig.2E***), our findings establish that induced chemokine production is independent of mouse strain, immunodominant determinants and functional avidities but quantitatively different in specific CD8^+^ and CD4^+^T^E^.

To provide a rough estimate for the relative contribution of chemokine production to the totality of quantifiable CD8^+^T^E^ functionalities, we determined the respective percentages of NP^396^-specific CD8^+^T^E^ capable of individual chemokine, cytokine and TNFSF ligand synthesis; constitutive GzmA/B and perforin expression; and degranulation/killing; according to this estimate, >40% of the CD8^+^T^E^ response is in fact dedicated to chemokine production (***Fig.2F***).

### Similar chemokine expression profiles of LCMV-, VSV-, LM- and vaccine-specific CD8^+^ and CD4^+^T^E^

The regulation of pathogen-specific CD8^+^ and CD4^+^T immunity generated in response to vesicular stomatitis virus (VSV) or LM shares many cardinal properties with the LCMV system [45–51] yet the distinct biology of these pathogens may have an impact on aspects of the T cell chemokine response. In contrast to the non-cytopathic arenavirus and natural murine pathogen LCMV, VSV is an abortively replicating cytopathic virus that causes a polio- or rabies-like neurotropic infection in immunodeficient mice [52]. Similar to other pathogenic bacteria such as Mycobacteria, Salmonella, Rickettsia and Chlamydia, LM is a facultative intracellular bacterium and the model of murine listeriosis constitutes one of the best-characterized experimental systems for bacterial infection [53]. Acute infection with LM (rLM-OVA for induction of a traceable specific CD8^+^T^E^ population) or VSV generated specific CD8^+^T^E^ with constitutive CCL5 and inducible CCL3/4/5/9/10 and XCL1 expression akin to those found in LCMV-specific CD8^+^T^E^; similarly, LM- and VSV-specific CD4^+^T^E^ presented with chemokine production profile resembling LCMV-specific CD4^+^T^E^ (***Figs.3A/B & S3D***). The considerable uniformity of chemokine signatures by T cells specific for three disparate pathogens therefore identifies fundamental functional attributes of the pathogen-specific T cell response at large (***Figs.2E & 3C***).

**Figure 3.**
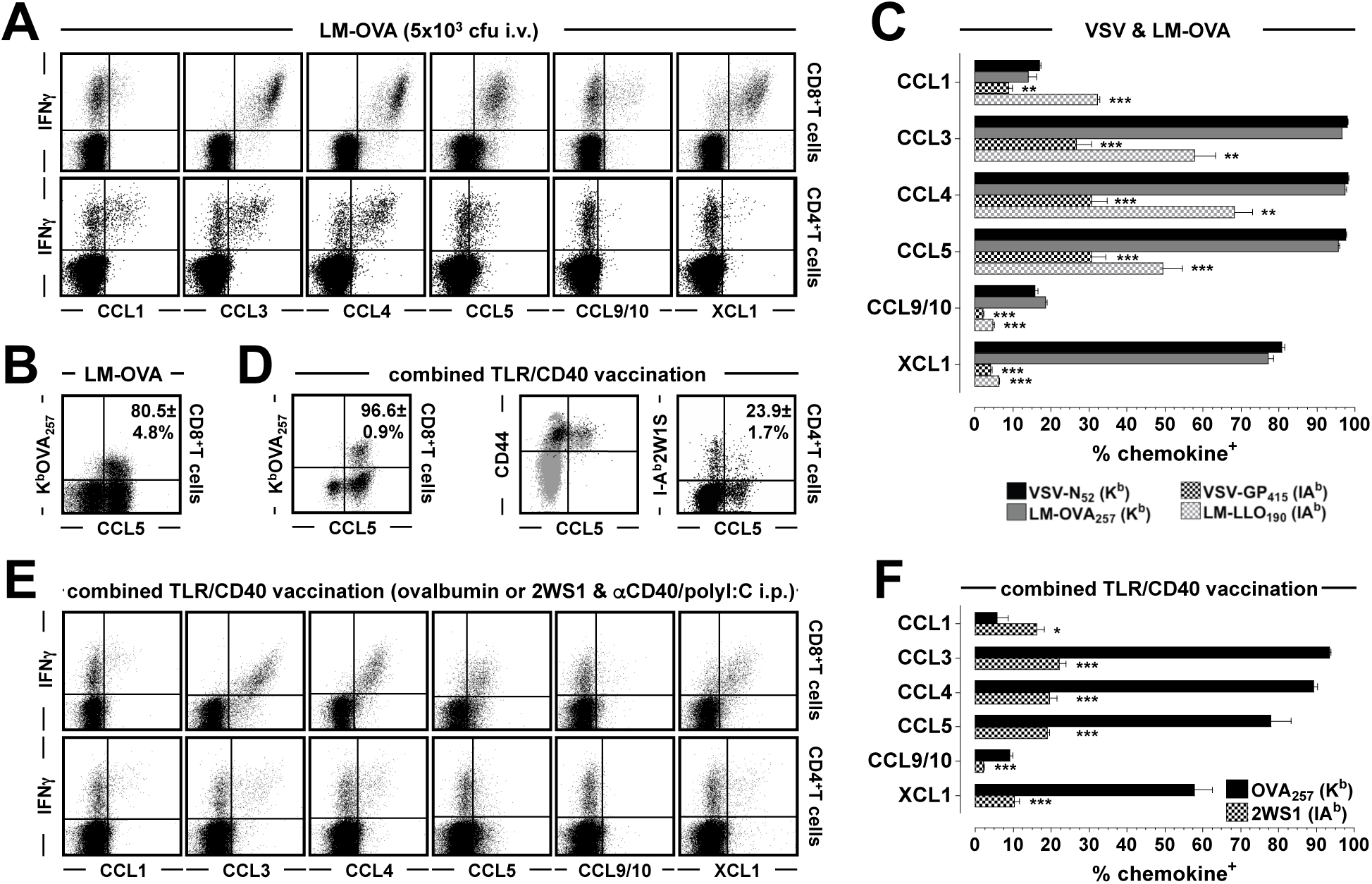
Constitutive and induced chemokine expression by LM-, VSV-, and vaccine-specific CD8^+^ and CD4^+^T_E_. **A.,** induced chemokine expression by specific CD8^+^ and CD4^+^T_E_ (d8) following challenge with rLM-OVA (data display as in Fig.2C). **B.,** constitutive CCL5 expression by rLM-OVA_257_-specific CD8^+^T_E_ (d8). **C.,** summary of induced chemokine production by rLM-OVA- and VSV-specific CD8^+^ and CD4^+^T_E_ (restriction elements are indicated); asterisks denote significant differences between CD8^+^ and CD4^+^T_E_ specific for the same pathogen. **D.,** *ex vivo* CCL5 expression by vaccine-specific CD8^+^ and CD4^+^T_E_ (d7 after vaccination as explained below); dot plots are gated on blood-borne CD8^+^ or CD4^+^T cells as indicated, the two-tone dot plot is gated on both total CD4^+^T cells (gray) and I-A^b^2WS1^+^ CD4^+^T_E_ (black). **E./F.,** induced chemokine profiles of specific CD8^+^ and CD4^+^T_E_ generated by combined TLR/CD40 vaccination. Mice were challenged with ovalbumin/αCD40/polyI:C (“CD8^+^ vaccination”) or 2WS1 peptide/αCD40/polyI:C (“CD4^+^ vaccination”) as detailed in Methods and analyzed 7 days later; asterisks indicate differences between CD8^+^ and CD4^+^T_E_. All data (SEM) are representative for multiple experiments comprising groups of 3-5 individual mice.

Nevertheless, we noted some quantitative differences associated with the use of different infection protocols, and to ascertain if the degree of infection-associated inflammation could modulate T^E^ chemokine production profiles in a given model system, we infected B6 mice with escalating dosages of rLM-OVA (3×10^2^-3×0^4^ cfu). As expected, an increase of bacterial dosage heightened early inflammation as determined by serum IFNγ levels, but the numbers of OVA^257^-specific CD8^+^T^E^ as well as their *ex vivo* CCL5 expression levels peaked after infection with 3×10^3^ cfu rLM-OVA and declined at higher infection dosages; in contrast, LLO^190^-specific CD4^+^T^E^ numbers steadily rose with escalating challenge dosages (***Fig.S4A***). While the fraction of IL-2-, TNFα- and CD40L-producing specific CD8^+^ and CD4^+^T^E^ remained impervious to bacterial challenge dosage, induced chemokine synthesis by OVA^257^-specific CD8^+^T^E^ was compromised by infection with higher rLM-OVA titers, especially the production of CCL1 and XCL1 and to a lesser extent also CCL3/4/5; the chemokine expression profiles of LLO^190^-specific CD4^+^T^E^, however, remained largely unaffected by the different challenge protocols (***Fig.S4B/C***). These observations suggest that chemokine production by CD8^+^ but not CD4^+^T^E^ may be partially impaired under conditions of chronic infection, and we have further pursued this question in related work (manuscript in preparation).

We also extended our delineation of chemokine expression profiles to vaccine-specific CD8^+^ and CD4^+^T^E^. Using a strategy for induction of protective T cell immunity by combined TLR/CD40 vaccination, *i.e.* the immunization with whole proteins or peptides in conjunction with poly(I:C) and αCD40 administration [54–56], we found that vaccine-induced CD8^+^ and CD4^+^T^E^ were remarkably similar to the respective pathogen-specific T^E^ populations at the level of constitutive (CCL5) and induced chemokine production capacity (***Fig3.D-F***). Both effective vaccines and different infectious pathogens therefore elicit essentially the same T^E^ chemokine response that is quantitatively adjusted according to the particular conditions of T cell priming.

Finally, small subsets of both LCMV- and LM-specific CD4^+^T^E^ have been described to exhibit a “T^H^2 phenotype” [57, 58]. While the existence of specific IL-4-producing CD4^+^T^E^ in these model sytems has been contested by others [50, 59], the description of CXCL2 production as a characteristic for *in vitro* generated T^H^2 cells [24, 60] permits an analysis of T^H^2 functionality at the chemokine level. Indeed, primary murine T cells expressed CXCL2 after polyclonal activation preferentially under exclusion of IFNγ (***Fig.S1A***), and a very small subset of LCMV-but not rLM-OVA-specific CD4^+^T_E_ produced CXCL2 (***Fig.S3E***). However, given the clearly predominant “T_H_1 phenotype” of LCMV- and LM-specific CD4^+^T cells [47, 50, 59], we have not pursued a further characterization of “T_H_2 chemokines” in these model systems.

### Delayed acquisition of CCL5 production capacity by CD8^+^T_E_

The acquisition of defined effector functions constitutes a hallmark of primary T_E_ differentiation and the detailed work by A. Krensky’s group has identified an unusual property of T cell-produced CCL5, namely it’s comparatively late synthesis only after 3-5 days of TCR stimulation as a consequence of regulation through the transcription factor KLF13 [61–63]. To elucidate these dynamics for pathogen-specific T cells, we compared the regulation of CCL5 expression with that of principal effector molecules (GzmB and IFNγ) during the transition from naïve to early effector stage of developing p14 T_E_. Assayed over a 72h period *in vitro*, the rapid and progressive induction of IFNγ and slightly delayed GzmB synthesis as a function of cell division contrasted with a lack of constitutive and only minimal inducible CCL5 expression (***Fig.4A/B***). Similarly, within the first 60h after *in vivo* challenge, p14 T_E_ remained CCL5-negative and constitutive CCL5 expression by ∼50% of p14 T_E_ or endogenously generated CD8^+^T_E_ became discernible only on d5 after LCMV infection; by d8, however, practically all LCMV-specific CD8^+^T_E_ had acquired a CCL5^+^/GzmB^+^ phenotype (***Fig.4C/D***). The protracted dynamics of CCL5 expression are indeed unique as they differed not only from GzmB and IFNγ but also GzmA and all other inducible CD8^+^T_E_ functionalities evaluated (CCL1/3/4, XCL1, IL-2, GM-SF, TNFα and CD40L; not shown); accordingly, constitutive CCL5 protein expression may serve as a novel functional marker for mature antigen-experienced CD8^+^T_E_.

**Figure 4.**
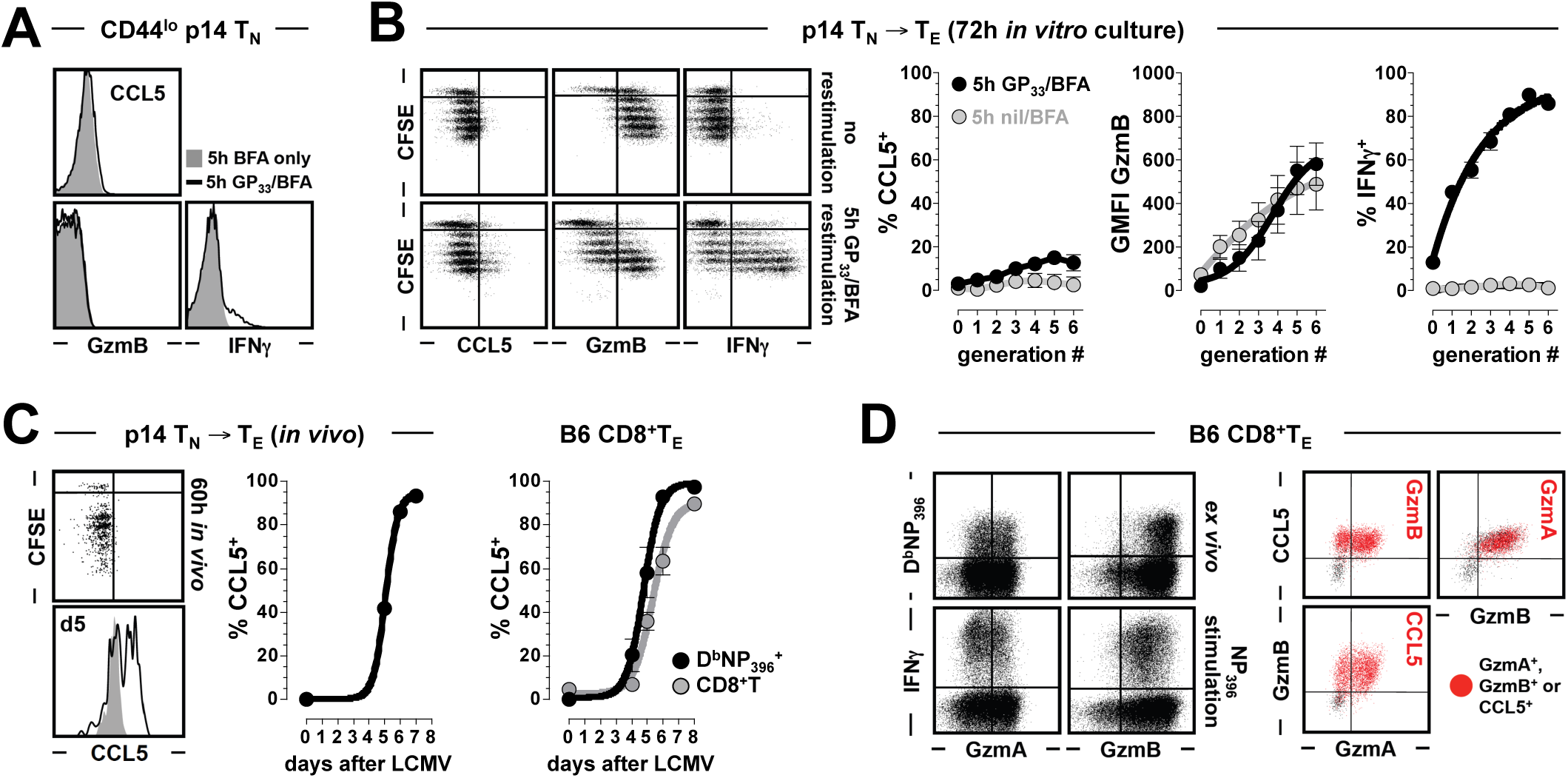
Expression and acquisition kinetics of CD8^+^T_E_ effector molecules. **A.,** constitutive (gray histograms) and induced (black tracings) CCL5, GzmB and IFNγ expression levels by naïve CD44^lo^ p14 cells (p14 T_N_; note that the functionality of p14 T_N_ is restricted to limited IFNγ production). **B.,** CCL5, GzmB and IFNγ expression as a function of early *in vitro* p14 T_E_ proliferation. Dot plots are gated on p14 T cells analyzed directly after 72h stimulation culture (“no restimulation”) or following GP_33_ peptide restimulation in the presence of BFA during the final 5h of culture as indicated. The diagrams on the right summarize the individual expression patterns as a function of p14 CFSE dilution (generation #0: no division). **C.,** acquisition of constitutive CCL5 expression by p14 CD8^+^T_E_ *in vivo* was analyzed after 60h after adoptive transfer of CFSE-labeled p14 cells and LCMV challenge (top dot plot) or in p14 chimeras on day 5 after infection (bottom plot gated on blood-borne CCL5^-/-^ [gray] and wt [black] p14 T_E_); the adjacent diagrams depict the emergence of constitutive CCL5 expression by developing CD8^+^T_E_ in the p14 chimera system (middle) and LCMV-infected B6 mice (right diagram: fraction of CCL5^+^ T cells among total [gray] and D^b^NP_396_^+^ [black] CD8^+^T cells). **D.,** left: GzmA and GzmB expression by specific CD8^+^T_E_ (d8) analyzed *ex vivo* (top) or after 5h restimulation culture (bottom); all dot plots gated on splenic CD8^+^T cells. Right: constitutive CCL5, GzmA and GzmB expression by LCMV-specific CD8^+^T_E_ (the small subset of CCL5- and GzmA/B-negative CD8^+^T cell subset corresponds to the CD44^lo^ naïve CD8^+^T cell fraction, not shown).

### Constitutive co-expression and subcellular localization of CCL5 and granzymes in antiviral CD8^+^T_E_

The precise subcellular localization of CCL5 remains a matter of controversy. In humans, a reported preferential association with the content of cytolytic granules (GzmA, perforin, granulysin) [17, 64] contrasts with the identification of a unique subcelluar CCL5 compartment [65], and the frequent use of T cell clones or blasts, the differential regulation of cytolytic effector gene and protein expression in primary murine CTL [44, 66–68], and the previously reported absence of constitutive CCL5 expression by mouse CD8^+^T_MP_ in particular [69, 70] further complicate resolution of this issue. Our direct *ex vivo* FC analyses of LCMV-specific CD8^+^T_E_ now demonstrate a clear association of CCL5 and GzmB expression while GzmA, as reported previously by us (and also similar to influenza-specific CD8^+^T_E_ [34, 44]), was expressed by only ∼60% of the CD8^+^T_E_ population (***Fig.4D***). These observations indicate that CCL5 co-localization studies in murine CD8^+^T_E_ should be extended beyond the visualization of GzmA. Accordingly, confocal microscopy revealed the existence of multiple discrete vesicles that contained either GzmA, GzmB or both (***Fig.5*** rows 1 & 2); in contrast, CCL5^+^ vesicles appeared mostly devoid of GzmA/B (***Fig.5***, rows 3-5) and thus presented with an expression patterns reminiscent of the subcellular CCL5 localization in primary human CD8^+^T_MP_ [65]. Nevertheless, we did observe polarization and coalescence of GzmA/B^+^ and CCL5^+^ vesicles in some cells, perhaps a result of recent CD8^+^T_E_ activation and an indication that these effector molecules are likely co-secreted (***Fig.5***, row 6 & 7). The overall distribution of granzymes and CCL5 across individual vesicles in CD8^+^T_E_ is therefore somewhat heterogeneous, a conclusion also reported for the subcellular expression patterns of CCL5, perforin and granulysin in human CD8^+^T cells [64].

**Figure 5.**
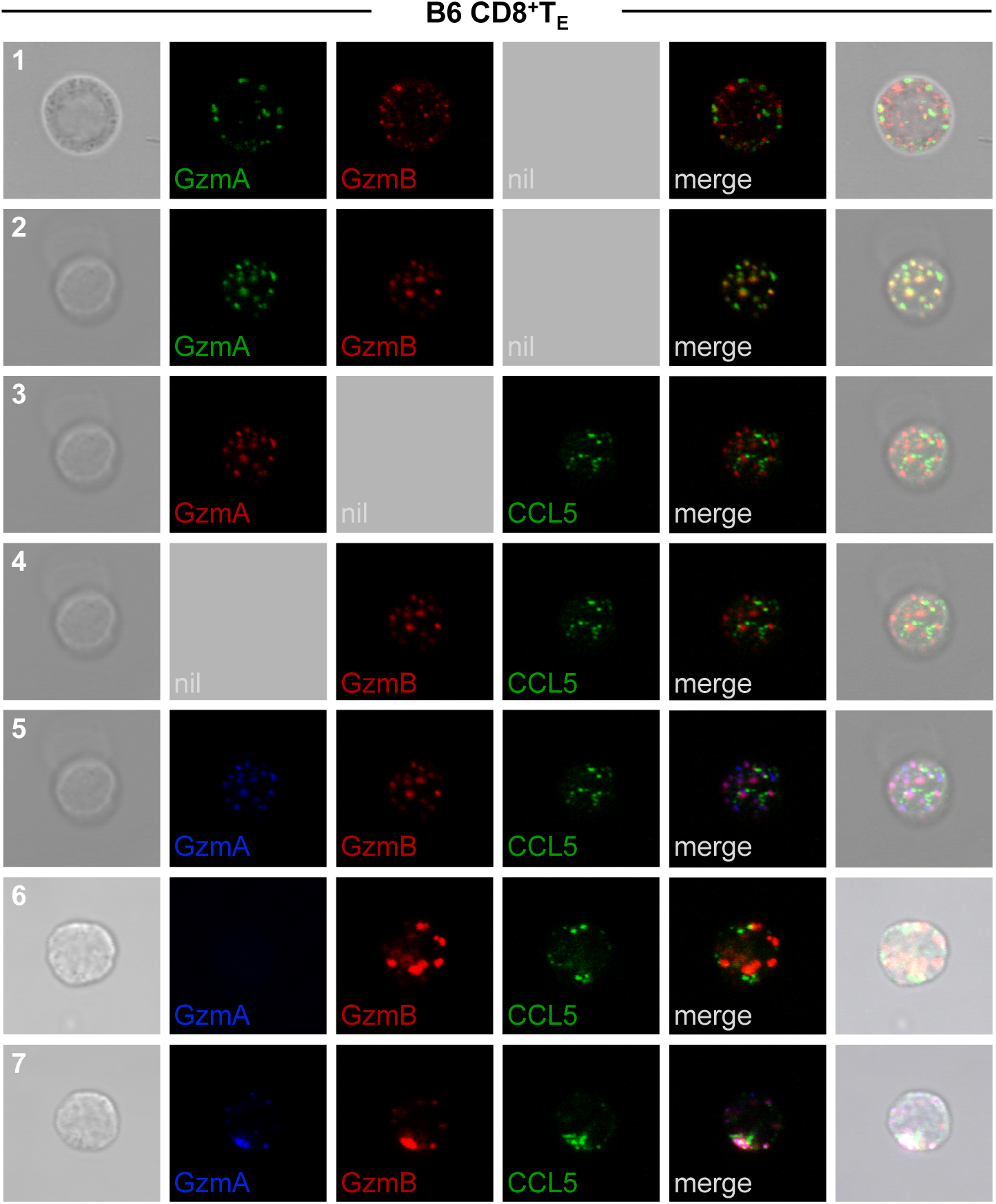
Subcellular localization of GzmA, GzmB and CCL5 in CD8^+^T_E_. Blood-borne CD8^+^T_E_ (d8) were stained with αGzmA-FITC, αGzmB-APC and αCCL5/αgoat-Cy3, and sorted GzmA^+^ subsets were analyzed by confocal microscopy as detailed in Methods (GzmA^-^ subsets were used as a negative staining control). Rows 1 & 2: subcellular GzmA and GzmB localization in 2 different cells; rows 2-5: same cell analyzed for GzmA, GzmB and/or CCL5 co-localization; row 6: GzmB and CCL5 expression in the sorted GzmA^-^ subset; row 7: example for partial polarization and coalescence of intracellular GzmA/B and CCL5 stores in another CD8^+^T_E_.

The precise molecular mechanisms underpinning constitutive CCL5 expression by CD8^+^T_E_, unique among the cytokines/chemokines, remain unclear. Given the general similarity between CD8^+^T and NK cells [71], and more specifically the chemokine production profiles largely shared between CD8^+^T_E_ and NK cells [33], we considered the proposal that constitutive CCL5 expression by human NK cells is dependent on the JNK pathway [72]. As shown in ***Fig.S4D/E***, however, Jnk1^-/-^ and Jnk2^-/-^ mice, both of which readily control an LCMV infection [73], did not present with any abnormalities at the level of constitutively expressed CCL5 in NK cells or virus-specific CD8^+^ or CD4^+^T_E_.

### Kinetics of chemokine synthesis by virus-specific CD8^+^T_E_

The elaboration of diverse T cell effector functions is a coordinated event that integrates spatial and temporal constraints with potentially different activation requirements. To determine the velocity of chemokine production by specific CD8^+^T_E_, p14 T_E_ were restimulated for 0-5h with GP_33_ peptide in the presence or absence of transcriptional (actinomycin D: ActD), translational (cycloheximide: CHX) and/or protein secretion inhibitors (brefeldin A: BFA), and analyzed for intracellular chemokine content by FC using IFNγ production as a reference. Induced IFNγ, CCL3 and CCL4 expression became detectable after as little as 30min of stimulation and reached a maximum after 4-5h. The synthesis of these proteins was sufficiently robust to allow detection of intracellular IFNγ and CCL3/4 even in the absence of BFA (***Fig.6A-C***, panels 1 & 2). Given the presence of *ex vivo* detectable mRNA species for CCL3/4 and IFNγ (***Figs.1A & S3A***), protein synthesis, while reduced, was still observed under conditions of transcriptional blockade in the presence but not absence of BFA (***Fig.6A-C***, panels 3 & 4). The fact that protein synthesis increased over time in a homogenous fashion in all p14 T_E_ (not shown) suggests that constitutive IFNγ and CCL3/4 message is evenly distributed among individual cells rather than preferentially allocated to a particular p14 subset. As expected, inhibition of translation or combined transcriptional/translational blockade completely prevented the accumulation of IFNγ a□□□CCL3/4 proteins (***Fig.6A-C***, panels 5-8).

**Figure 6.**
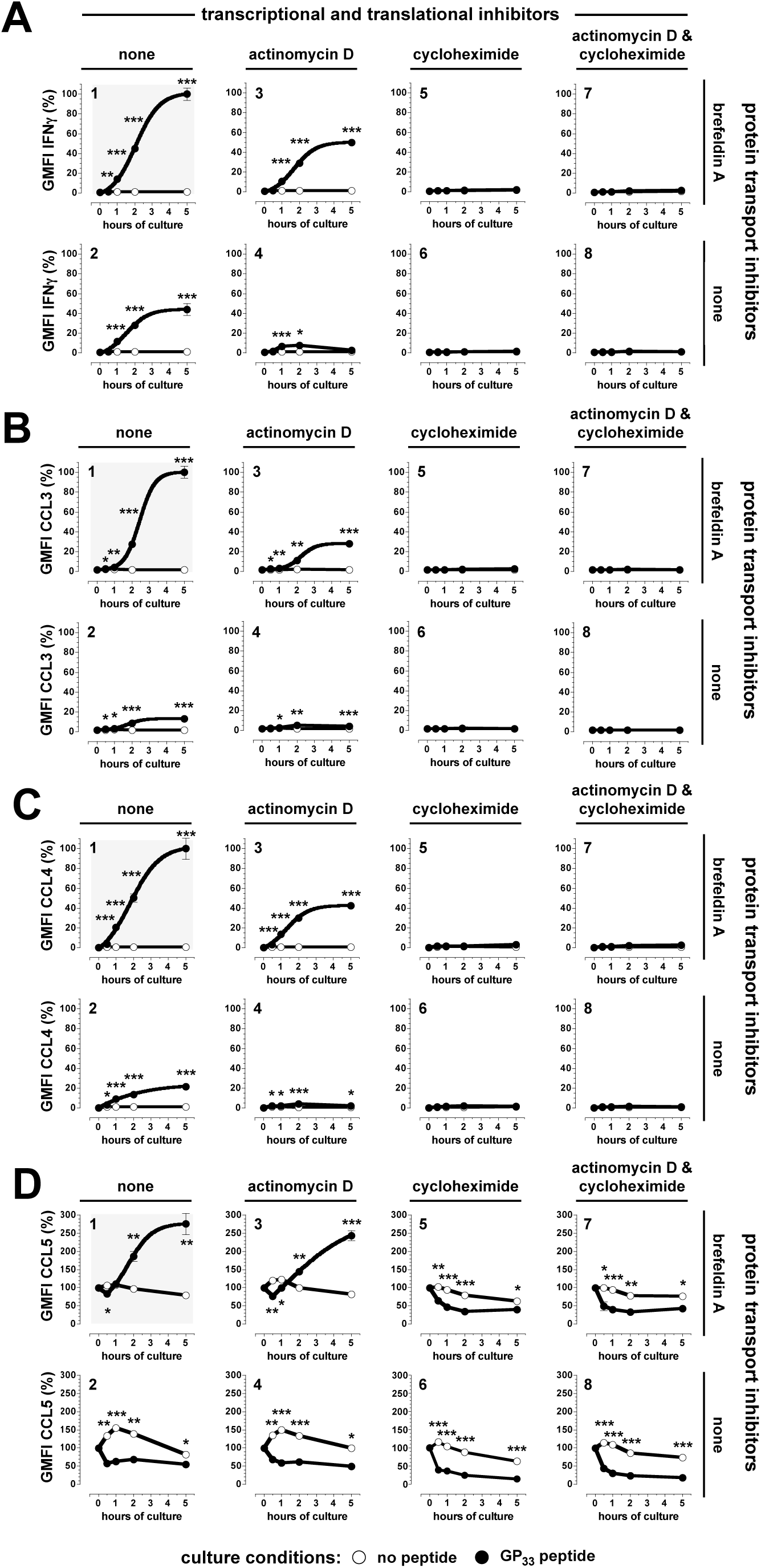
Regulation and kinetics of chemokine production by p14 CD8^+^T_E_. **A.-D.,** spleen cells from LCMV-infected p14 chimeras (d8) were cultured for indicated time periods in the presence (closed symbols) or absence (open symbols) of GP_33_ peptide and indicated transcriptional (ActD), translational (CHX) and/or protein transport (BFA) inhibitors. Graphs depict the GMFI of IFNγ, CCL3, CCL4 and CCL5 expression by p14 T_E_ as a function of culture time and inhibitor presence or absence (panels depicting “traditional” stimulation conditions, *i.e.* peptide plus BFA, are shaded in gray). To better compare the kinetic regulation of cytokine and chemokine production, the respective GMFI values were normalized (IFNγ: the GMFI of rat isotype control stains was subtracted from all corresponding IFNγ GMFI values and the resulting values of BFA/GP_33_ cultures for the t=5.0h time point (panel A.1) were set at 100%. CCL3 and CCL4: similar normalization performed by subtraction GMFI values of goat IgG stains from corresponding CCL3 or CCL4 GMFI values. CCL5: *ex vivo* goat IgG control stain GMFI was subtracted from all CCL5 GMFI values and resulting normalized *ex vivo* CCL5 values (panel B.1) were set at 100%). The sigmoidal curve fit is based on optimal fits determined by non-linear regression analyses of samples containing additional time points (n=3 mice/group, data from 3 similar experiments).

The kinetics of intracellular CCL5 accumulation were predictably more complex since TCR-induced release of pre-stored CCL5 and initiation of protein neosynthesis occurred in parallel. In fact, a modest loss of CCL5 observed 30min after TCR stimulation was quickly compensated by a pronounced increase of intracellular CCL5 in cultures containing BFA, a pattern that contrasted with rapid if only partial CCL5 depletion in the absence of BFA (***Fig.6D***, panels 1 & 2). It is therefore worth mentioning that the release of newly synthesized CCL5, in contrast to the release of pre-stored CCL5 [65], was largely inhibited by BFA. Furthermore, the kinetics of intracellular CCL5 accumulation were comparable in the presence and absence of transcriptional inhibition (***Fig.6D***, panels 1 & 3) consistent with our observation that TCR stimulation does not increase the level of CCL5 mRNA (***Fig.1A***), and an early increase of intracellular CCL5 in cultures without TCR stimulation and translational blockade emphasizes that maintenance of constitutive CCL5 expression by T cells is an active process; the eventual decline of CCL5 expression at later time points is likely due to degradation since we did not observe CCL5 secretion by unstimulated T cells (***Fig.6D***, panels 2 & 4 and not shown). Interestingly, upon TCR stimulation in the presence of translational or combined transcriptional/translational blockade, ∼2/3 of pre-stored CCL5 were released within 30-60min; additional depletion of CCL5 stores occurred with slower kinetics and was inhibited by BFA (***Fig.6D***, panels 5-8 ***& Fig.7A***).

**Figure 7.**
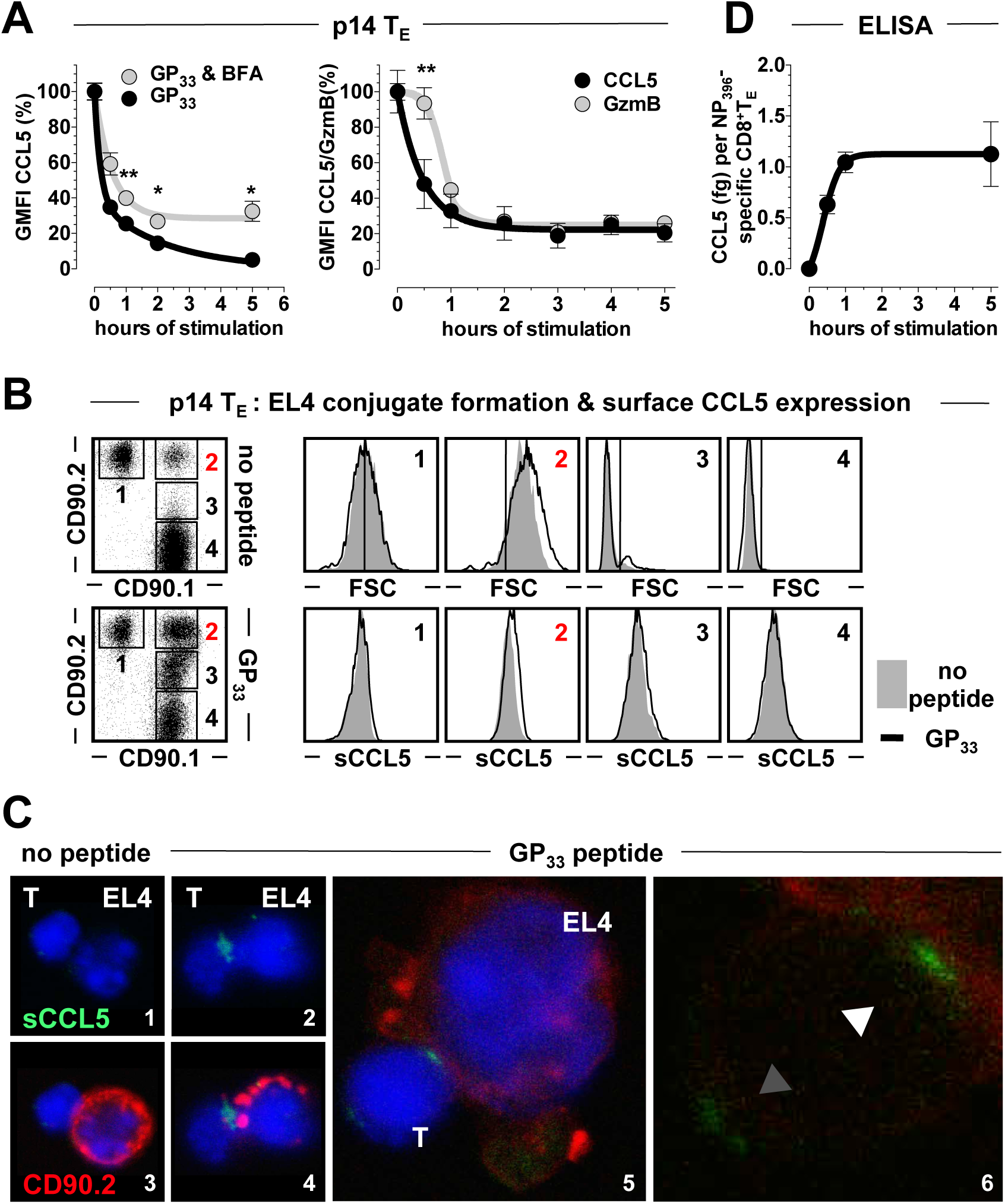
Rapid surface translocation and secretion of pre-stored CCL5 by antiviral CD8^+^T_E_. **A.,** spleen cells from d8 p14 chimeras were pre-incubated with CHX prior to initiation of TCR stimulation by addition of GP_33_ peptide and subsequent analysis of p14 GzmB and CCL5 content 0-5h later. GzmB and CCL5 expression levels (GMFI) were normalized such that t=0h levels correspond to 100% and the GMFI of respective control stains are set at to 0%. Left: kinetics of pre-stored CCL5 release in the presence *vs.* absence of BFA. Right: “immediate” depletion of CCL5 stores *vs.* delayed GzmB release. At t=0.5h, ∼2/3 of CCL5 but <10% of GzmB stores are emptied (n=3 mice, 1/3 independent experiments). **B.,** conjugate formation between purified p14 T_E_ (CD90.1) and GP_33_ peptide-coated or uncoated EL4 cells (CD90.2) as well as CCL5 surface expression (sCCL5) were assessed 20min after initiation of co-culture as detailed in Methods. Four populations were distinguished according to CD90.1/2 expression levels and FSC properties (forward scatter, cell size): 1., EL4 cells; 2., EL4:p14 T_E_ conjugates; 3., p14 T_E_ expressing low levels of CD90.2, likely acquired by trogocytosis; and 4., p14 T_E_. Note the weak but distinctive sCCL5 staining detectable among specific (black tracings) but not unspecific (gray histograms) EL4:p14 T_E_ conjugates (population 2). **C.,** conjugation assays were performed as above and analyzed by confocal microscopy to visualize sCCL5 (green) and CD90.2 (red) expression (panels 1/3 and 2/4 are identical with CD90.2 signals removed from panels 1 and 2 to better visualize sCCL5 expression). Note the “blebbing” of the EL4 cell in panel 5 consistent with the induction of apoptosis; panel 6 features a magnification of the p14 T_E_ in panel 5 to demonstrate IS (white arrow) and antipolar (gray arrow) localization of sCCL5. **D.,** spleen cells from LCMV-infected B6 mice (d8) were pre-incubated with CHX, stimulated with NP_396_ peptide (no BFA) and CCL5 in the supernatant quantified by ELISA. To calculate the amount of pre-stored CCL5 secreted by individual NP_396_-specific CD8^+^T_E_, complementary FACS analyses were performed to determine the absolute numbers of cultured specific CD8^+^T_E_. Further, the amount of CCL5 secreted in the absence of TCR stimulation was subtracted from stimulated samples at all time points. Note that after 30min, ∼60% of total CCL5 is already secreted, at 1h, ∼90%.

### Rapid CCL5 surface translocation and secretion by virus-specific CD8^+^T_E_

To interrogate the remarkably fast CCL5 release kinetics in more detail, we compared the concurrent depletion of pre-stored CCL5 and GzmB from TCR-stimulated p14 T_E_ in the presence of CHX. Here, the near instantaneous release of CCL5 contrasted with a ∼30min lag period before intracellular GzmB began to decline. Yet the subsequent loss of GzmB proceeded so rapidly that the relative extent of CCL5 and GzmB depletion was comparable by 1h after initiation of T cell stimulation (***Fig.7A***). Overall, the kinetic differences between CCL5 and GzmB depletion as well as the differential sensitivity of constitutive *vs.* induced CCL5 expression/release to BFA corresponds well with the heterogeneous distribution of CCL5 and GzmB across different subcellular compartments as shown in ***Fig.5***.

In order to better visualize the earliest events of TCR-induced CCL5 release, we performed an *in vitro* conjugation assay using purified p14 T_E_ (CD90.1) and congenic (CD90.2) GP_33_ peptide-coated *vs.* uncoated EL4 target cells (***Fig.7B/C***). Within 20min after TCR engagement and thus before initiation of CCL5 neosynthesis, CCL5 was translocated to the cell surface, deposited preferentially at the interface between p14 T_E_:EL4 conjugates, and the engagement of EL4 cells was readily demonstrated by the focused redistribution of CD90.2 around the immunological synapse (IS) [74]; formation of “unspecific” conjugates (*i.e.,* in the absence of GP_33_ peptide) was not associated with cell surface exposure of CCL5 nor a clustering of EL4-expressed CD90.2 (***Fig.7B/C***). Our data therefore support the notion that mobilization of intracellular CCL5 stores is primarily directed towards the IS similar to the polarization reported for polyclonally activated human CD8^+^T cell blasts and clones [65, 75]; yet they apparently differ from results of another study in which the *de novo* synthesis of CCL5 by *in vitro* generated murine CD4^+^T cell blasts resulted in multidirectional release of this chemokine, *i.e.* an early (2h) association of intracellular chemokine stores with the IS followed by a later (4h) distribution in multiple compartments throughout the cytoplasm [76]. In agreement with the latter report, however, we found that p14 T_E_ on occasion presented low amounts of antipolar surface CCL5 (***Fig.7C***, panel 6) and it is tempting to speculate that the heterogenous subcellular distribution of pre-stored CCL5 is related to the reported association with distinct trafficking proteins that mediate a multidirectional *vs.* focused chemokine release [75, 76].

The rapid accumulation of CCL5 within the IS, a defined space with an estimated volume of 0.5-5.0×10^-16^ liters [76, 77], suggests that local CCL5 concentrations may temporarily reach “supra-physiological” levels. The latter term describes multiple observations that *in vitro* exposure of cells to oligomeric CCL5 in excess of ∼1μM promotes receptor-independent binding to surface glycosaminoglycans and generalized activation that, depending on the cell type under study, results in cellular proliferation and differentiation as well as enhanced survival, CTL activity, cytokine and chemokine release [78–84]. If these effects have an *in vivo* correlate has remained doubtful and to provide a more quantitative estimate, we combined FC and ELISA assays conducted in the presence of CHX to calculate the amount of pre-stored CCL5 that is secreted by an individual LCMV-specific CD8^+^T_E_: the rapid increase of CCL5 in the ELISA culture supernatant (***Fig.7D***) mirrors the loss of intracellular CCL5 in ***Fig.7A*** and corresponds to ∼0.5fg per CD8^+^T_E_ released in the first 30min after TCR triggering, an amount that could in principle result in a CCL5 concentration of >100μM within the confines of the IS. Even if these calculations constitute a gross overestimate due to incomplete CCL5 release, multiple CD8^+^T_E_:target cell contact sites [85], limited spatial constraints and/or rapid diffusion, it would appear likely that CCL5 concentrations of >1μM could be achieved in a spatially and temporally confined fashion *in vivo* and therefore might contribute to target cell activation in a receptor-independent fashion.

### Induced CCL3/4/5 co-localization and co-secretion by virus-specific CD8^+^T_E_

The co-production of CCL3/4/5 by stimulated CD8^+^T_E_, as evidenced by the “diagonal” event distribution in FC plots (***Fig.2D***), suggests a tight association and potential co-localization of these chemokines following CD8^+^T_E_ activation. When analyzed by confocal microscopy under these conditions, CCL3/4/5 as well as GzmB and IFNγ indeed tended to cluster in a single defined location close to the plasma membrane and in immediate proximity of the IS (***Fig.S5A*** and not shown). The co-localization of induced CCL3/4/5 expression in particular might provide a basis for the joint release of these chemokines bound to sulphated proteoglycans as described for HIV-specific CD8^+^T cell clones [17]. Moreover, human PBMC stimulated with PMA secrete CCL3/4 as heterodimeric complexes [86] and up to half of the CCL3/4 content in medium conditioned with LN cells from recently immunized mice is in a state of hetero-oligomerization [29]. To determine the extent of heterologous chemokine complex formation in the context of a virus-specific T cell response, splenic CD8^+^T_E_ were restimulated for 5h, the supernatants collected and pre-absorbed with αCCL3, αCCL4, αCCL5 or control antibodies prior to quantitation of CCL3/4/5 by ELISA (***Fig.S5B***). Although we noted some variability in these cross-absorption experiments, we have previously confirmed the specificity of antibodies used for pre-absorption [33] and therefore can conclude that the biologically active form of CD8^+^T_E_-secreted CCL3/4/5 consists in part of hetero-oligomeric complexes. Beyond the apparently intimate coordination of CCL3/4/5 activities, this finding also emphasizes important limitations for the interpretation of any experiments that employ antibody-mediated *in vivo* neutralization of these chemokines.

### Specific antiviral T cell immunity in the absence of systemic chemokine deficiencies

Despite their prominence among T cell-produced effector molecules, CCL3/4/5 are apparently dispensable for the control of an acute LCMV infection [87–89]. Accordingly, we found that LCMV-challenged B6.CCL3^-/-^ and B6.CCL5^-/-^ mice generated a diversified virus-specific T cell response and controlled the infection with kinetics comparable to B6 wt mice (***Fig.S6A/B***; similarly, B6.CCL1^-/-^ mice mounted normal T_E_ responses and readily controlled LCMV, ***Fig.2D*** and not shown). Since CCL5 may exert direct apoptotic functions [90] and *in vitro* degranulation and killing by CCL5^-/-^ CD8^+^T cells in the context of a chronic LCMV infection is reportedly impaired [89], we also examined the *in vivo* killing kinetics by LCMV-specific CD8^+^T_E_ in the absence of CCL5. As shown in ***Fig.S6C***, however, *in vivo* target cell killing proceeded with the same rapid kinetics in wt and CCL5-deficient mice, and the lack of a role for CCL5 in this assay was further confirmed by treatment with a CCL5-neutralizing antibody.

Yet a careful analysis of T_E_ chemokine expression profiles in B6.CCL3^-/-^ and B6.CCL5^-/-^ mice demonstrated some unanticipated quantitative differences. In comparison to B6 mice, B6.CCL3^-/-^ but not B6.CCL5^-/-^ mice generated a slightly reduced antiviral CD8^+^ but increased CD4^+^T_E_ response (***Fig.S6A/B***). Furthermore, in B6.CCL3^-/-^ mice, CCL4 and CCL5 production by specific CD8^+^ and CD4^+^T_E_ was significantly diminished in comparison to B6 mice, and a somewhat lesser reduction of induced CCL3 and CCL4 expression was also observed for B6.CCL5^-/-^ mice (***Figs.8A/B & S6D***). These differences were even more pronounced when chemokine production by specific CD8^+^T_E_ was quantified *in vivo* following a 1h peptide inoculation (***Fig.8C*** and not shown). Lastly, both B6.CCL3^-/-^ and B6.CCL5^-/-^ mice also exhibited a modest but significant impairment of CCL9/10 production capacity by GP_33_-specific CD8^+^T_E_ interrogated *in vitro* (***Fig.S6D***). Altogether, functional impairments extending beyond specific chemokine gene deficiencies may be related to the cooperative regulation of chemokine expression/secretion and/or to artifacts arising in the mutant mice due to the proximity of respective gene loci on chromosome 11 (***Fig.1D***); they will also need to be considered for any interpretation of relevant observations made with CCL3- or CCL5-deficient mice,

**Figure 8.**
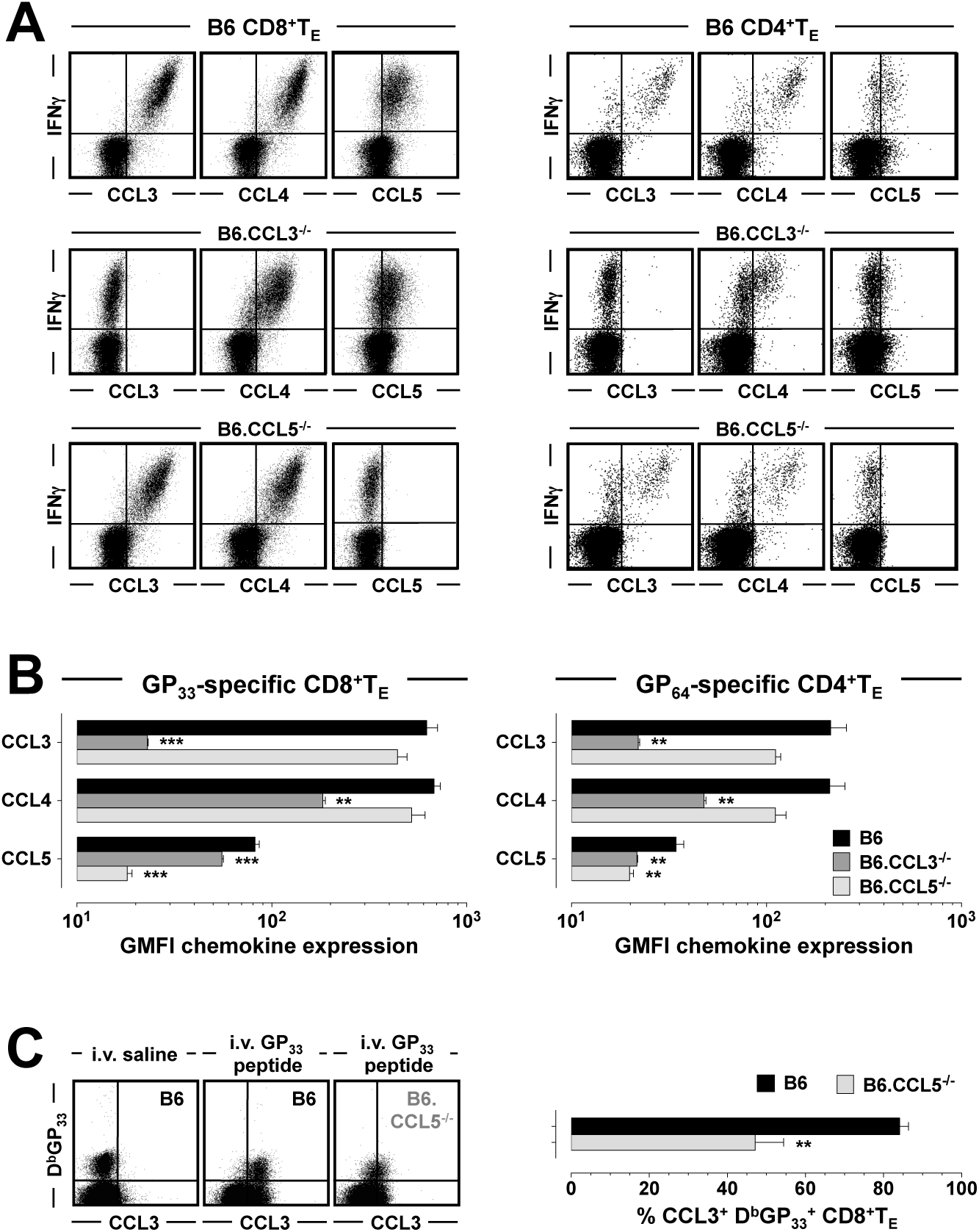
Impact of CCL3- or CCL5-deficiency on related chemokine production capacity by antiviral T_E_. **A.,** induced CCL3/4/5 production by GP_33_-specific CD8^+^ and GP_64_-specific CD4^+^T_E_ analyzed on d8 after LCMV challenge of B6, B6.CCL3^-/-^ and B6.CCL5^-/-^ mice (all plots gated on CD8^+^ or CD4^+^T cells). **B.,** CCL3/4/5 content of stimulated GP_33_-specific CD8^+^ and GP_64_-specific CD4^+^T_E_ in wt and chemokine-deficient mice (n=3/group, 1/3 similar experiments; astersisks indicate significant differences between B6 and mutant mice [one-way ANOVA]). **C.,** 1h *in vivo* CD8^+^T_E_ activation assays were performed on d8 after LCMV infection of B6 and B6.CCL5^-/-^ mice by i.v. injection of GP_33_ peptide as detailed in Methods (saline injection: negative control). Note the reduced CCL3 induction in CCL5-deficient D^b^GP_33_^+^CD8^+^T_E_ (n=3 mice/group; all plots gated on CD8^+^T cells; GP_33_ peptide also activates K^b^GP_34_^+^CD8^+^T_E_ accounting for the D^b^GP_33_ tetramer-negative population in the LR plot quadrants).

Lastly, fatal lymphocytic choriomeningitis following intracerebral (i.c.) LCMV infection of immunocompetent mice is contingent on a potent virus-specific CD8^+^T_E_ population that may recruit pathogenic myelomonocytic cells into the CNS through secretion of CCL3/4/5 [27, 91, 92]. Prior work with CCL3- and CCR5-deficient mice, however, has demonstrated a normal lethal phenotype after i.c. LCMV challenge [87, 88] leaving the possibility that CCL5 may uniquely contribute to the lethal disease. Here, we used a set of chemokine- and chemokine receptor-deficient mice to assess if the lack of any CD8T_E_-produced chemokines delayed or prevented lethal choriomeningitis. Specifically, we employed CCL1^-/-^, CCL3^-/-^, CCR5^-/-^ (CCL3/4/5 receptor), CCL5^-/-^, CCR1^-/-^ (CCL3/5/9/10 receptor) and CCR3^-/-^ (CCL5/9/10 receptor) mice and found that all of them succumbed to lethal disease with kinetics comparable to B6 or Balb/c control mice (***Fig.9***). Even CCL3-deficient mice lacking CCR5 and thus exhibiting a reduced CCL4/5 production capacity (***Fig.8A/B***) as well as decreased or absent responsiveness to CCL5 or CCL4, respectively, readily died after i.c. LCMV infection (***Fig.9***). While we cannot rule out that more complex compound chemokine/receptor deficiencies may alter the course of lethal disease and a potential contribution of the XCL1:XCR1 axis remains to be investigated, the fatal course of i.c. LCMV infection appears largely independent of chemokines produced by virus-specific CD8^+^T_E_.

**Figure 9.**
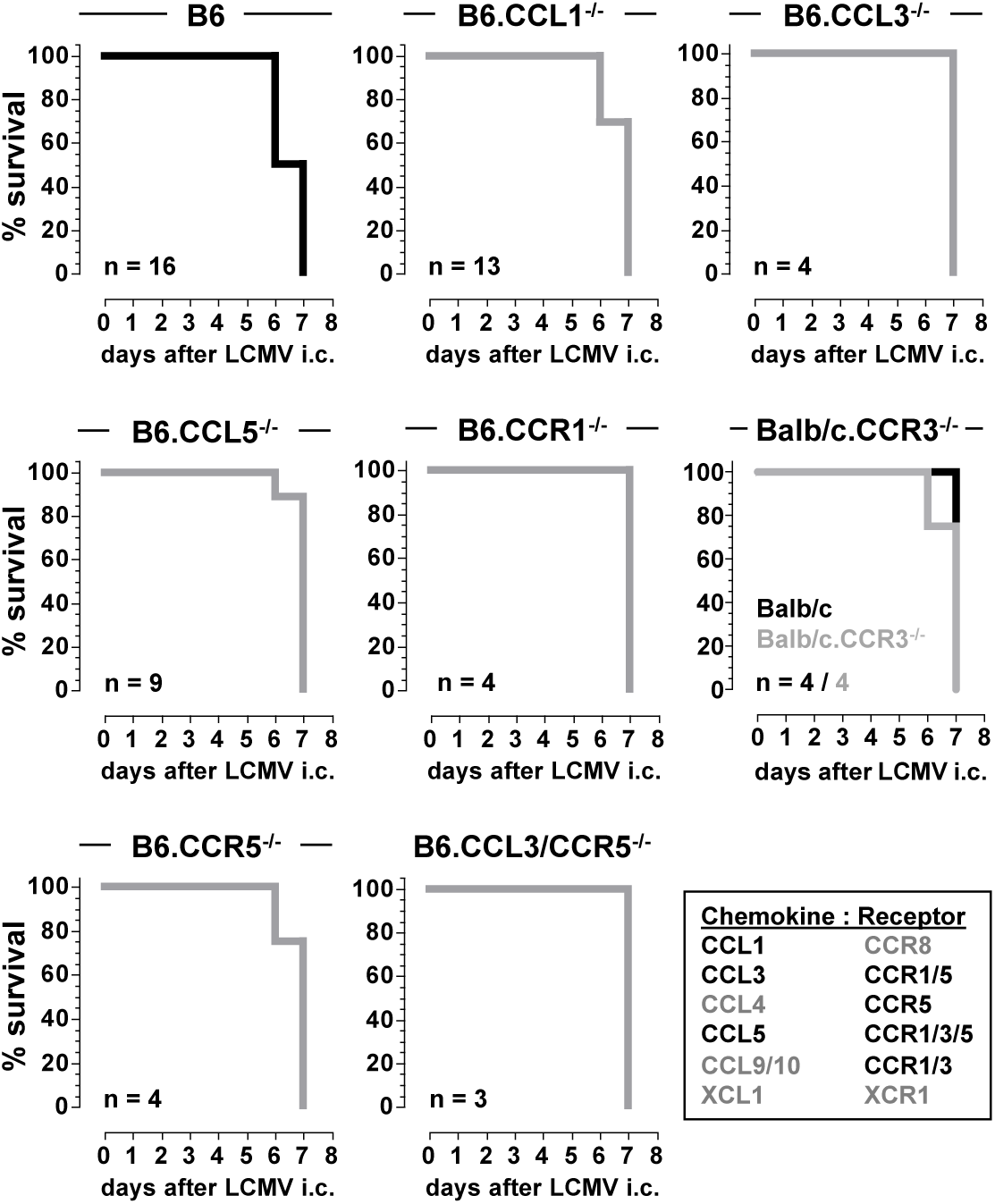
No role for antiviral T_E_-produced chemokines in the development of lethal choriomeningitis. Wild-type, chemokine- and/or chemokine receptor-deficient mice were infected with LCMV i.c. and survival was monitored (as per IACUC guidelines, we employed a scoring matrix to measure morbidity, and terminally ill mice were euthanized and scored as deceased). Multiple independent experiments were performed with matched experimental and control mice each, and the data displays feature the cumulative total (n) of individual mice analyzed. The lower right insert displays T_E_-produced chemokines and their respective receptors with specific chemokines/chemokine receptors interrogated in the present analysis highlighted in black.

## DISCUSSION

Pathogen-specific effector T cells are cardinal components of the adaptive immune response to viral and bacterial infections. Despite a wealth of knowledge about the contribution of T cell-derived cytokines, TNFSF ligands and cytolytic effector mechanisms to initial pathogen control, the full spectrum of potentially relevant T cell activities as well as their roles in shaping effective T_E_ responses and providing immune protection remain incompletely defined. Our delineation of the entire range of chemokines produced by specific CD8^+^ and CD4^+^T_E_ in the wake of different pathogen infections or immunizations constitutes an important addition to the analytically accessible repertoire of T cell functions for several important reasons: its near exclusive focus on six chemokines (CCL1, CCL3, CCL4, CCL5, CCL9/10, XCL1), a relative consistency of expression patterns across different *in vivo* challenge protocols, the sheer magnitude of the specific CD8^+^T_E_ chemokine response, and quantitative differences between CD8^+^ and CD4^+^T_E_ populations. In addition, T cells appear to be the major hematopoetic source for CCL1; CCL3/4/5 production/secretion is co-regulated; and the unique temporospatial organization of CCL5 synthesis, storage and secretion positions its targeted release at the forefront of the mature CD8^+^T_E_ response.

Using a combination of transcriptomic profiling and chemokine FC, we have defined a hierarchy of inducible chemokine expression by LCMV-specific CD8^+^T_E_ that pertains to all (CCL3/4) or nearly all (CCL5) CD8^+^T_E_ as well as greater and smaller subsets thereof (XCL1 > CCL1 ≥ CCL9/10) (constitutive CCL5 expression is discussed below). At the same time, no other chemokine proteins are synthesized by LCMV-specific CD8^+^T_E_, a contextual contention that is based on our use of a rigorously vetted collection of highly sensitive chemokine-specific antibodies deployed under optimal staining conditions [33]. While a lack of mRNA translation is a common feature of eukaryotic organisms [93], the absence of induced CXCL10 protein expression, in particular given the significant induction of *Cxcl10* mRNA following TCR stimulation, would appear to contradict reports that have documented CD8^+^T cell-produced CXCL10 (e.g., ref.[94]). To our knowledge, however, direct visualization of CD8^+^T cell-expressed CXCL10 has not been demonstrated (including in our own exploration of multiple other experimental scenarios), and CXCL10 detection in supernatants from T cell stimulation cultures can arise from small populations of contaminating myeloid cells that readily produce CXCL10 in response to T cell-secreted IFNγ (not shown). Although our conclusion that neither CXCL10 nor 30 other chemokines are produced by activated CD8^+^T_E_ is delimited by assay sensitivities and precise experimental context, the similarly restricted chemokine expression profiles of specific CD8^+^T_E_ across different epitope-specific populations with distinct avidities and immunodominant determinants, mouse strains and infection or vaccination modalities indicates that our analyses most likely capture the relevant components of the CD8^+^T_E_ chemokine response in their entirety. Thus, the inducible production of CCL1, CCL3, CCL4, CCL5, CCL9/10 and XCL1 is a shared signature of protective CD8^+^T_E_ populations generated in response to primary viral, bacterial and vaccine challenges. Moreover, the development of a vigorous chemokine response by vaccine-elicited CD8^+^T_E_, which in contrast to pathogen-specific CD8^+^T_E_ are not reliant on aerobic glycolysis to support their clonal expansion, reinforces the notion that the acquisition of robust effector functions is equally uncoupled from a “Warburg metabolism” [95]. It is also noteworthy that CCL9/10, regarded a “homeostatic” chemokine, is part of the “inflammatory” CD8^+^T_E_ response and therefore may be re-classified as a “dual function” chemokine; conversely, the constitutive expression of the “inflammatory” chemokines CCL3/4/5 by resting NK cells as shown here and/or in ref.[33] adds a “homeostatic” component that also may warrant the assignment of “dual function” to those chemokines.

One of the more striking aspects of the CD8^+^T_E_ chemokine response is its apparent magnitude. By stratifying CD8^+^T_E_ functionalities according to 17 individual parameters, we estimate that the synthesis and secretion of chemokines accounts for >40% of commonly quantified CD8^+^T_E_ activities. The remarkable abundance and distinct profile of chemokines produced by CD8^+^T_E_ therefore establish these cells as major focal points for the recruitment of other immune cells, the spatiotemporal organization of cellular interactions, and the overall coordination of complex effector immune responses. This conclusion, however, stands in marked contrast to the mostly modest phenotypes reported for specific T cell responses and/or pathogen control in mice lacking T cell-produced chemokines. Our own work confirms the generation of broadly normal LCMV-specific T_E_ responses and virus clearance kinetics in CCL3- and CCL5-deficient mice, and extends these observations to CCL1 deficiency; although these experiments do not specifically address the role of chemokines produced by T_E_, the lack of a pronounced phenotype under conditions of systemic chemokine deficiency strongly suggests a negligible function for the respective T_E_-derived chemokines. We further demonstrate in a stringent disease model where LCMV-specific CD8^+^T_E_ activities are essential for the recruitment of pathogenic myelomonocytic cells [27] that CCL1/3/4/5 and 9/10 are apparently dispensable for the development of lethal immunopathology. While these findings may in part be grounded in biological redundancies within the chemokine system, we note that lack of cardinal T_E_ molecules such as IFNγ, TNFα, IL-2, FasL, GzmA and/or GzmB often produce only subtle defects at the level of LCMV-specific T cell immunity and associated virus control [96–99]. Rather, non-redundant contributions of specific T cell-produced chemokines in effective control of primary pathogen infections are likely to emerge in the context of compound immune-deficiencies and within specific constraints of precisely delineated experimental scenarios for which the present study provides a comprehensive practical and conceptual foundation.

Another notable finding pertains to the chemokine response of CD4^+^T_E_ as well as its shared and distinctive aspects in comparison to CD8^+^T_E_ populations which typically present with substantially greater primary expansions [47]. The basic CD4^+^T_E_ chemokine profile, largely preserved in different infection and immunization settings, is composed of the same chemokines made by CD8^+^T_E_ (CCL1/3/4/5/9/10 and XCL1) but displays discrete quantitative differences: though CCL3/4/5 production is also the most prominent part of the CD4^+^T_E_ response, only 30-60% of pathogen-specific CD4^+^T_E_ readily synthesize these chemokines and the strict co-expression of CCL3/4 (not shown) points toward a specialized CD4^+^T_E_ subset dedicated to the chemokine-dependent recruitment of CCR1/3/5-bearing immune cells. The fraction of CCL1-producing CD4^+^T_E_ is comparable or somewhat larger than that of the corresponding CD8^+^T_E_ compartment, and only small subsets of CD4^+^T_E_ (≤ 5%) make CCL9/10 or XCL1. Interestingly, these pathogen-specific CD4^+^T_E_ chemokine profiles correspond remarkably well to a transcriptomic screen conducted for the presence of 28 chemokine mRNA species in *in vitro* polarized polyclonal T_H_1 cells [100]; the only other chemokine message detected in a complementary screen of T_H_2 cells was CXCL2 [100], also found at the protein level in polarized T_H_2 cells [24], and readily captured in our initial survey of primary T cell-produced chemokines. As expected for our “T_H_1-dominated” infection models, CXCL2-producing specific CD4^+^T_E_ were either absent (rLM-OVA) or present at very low frequencies (∼0.2%; LCMV).

Several of the T_E_-produced chemokines characterized here exhibit additional unique properties. T cells are considered an important source for CCL1 as evidenced, for example, by *Ccl1* mRNA transcription in activated CD4^+^ and CD8^+^T cell clones [101] or the secretion of CCL1 protein (in conjunction with the other CD4^+^T_E_ chemokines CCL3/4/5/9/10 and XCL1) by diabetogenic CD4^+^T cell clones [102]. Innate immune cells such as mast cells [103] and LM-infected DCs [104] may constitute additional hematopoetic sources for this chemokine but in our own work, we did not observe CCL1 expression by LM-infected DCs, activated B cells, myeloid cells or NK cells [33]; the lack of NK-cell-produced CCL1 is particularly noteworthy since these cells readily produce all of the other CD8^+^T_E_ chemokines [33]. Thus, while innate immune cell populations capable of CCL1 synthesis remain to be characterized in greater detail, T_E_ would appear to be a major and distinctive if not exclusive hematopoetic provenance of CCL1. Furthermore, its transcription/translation is strictly activation-dependent (i.e., *Ccl1* mRNA abundance displays the greatest differential of all T cell chemokines between *ex vivo* and αCD3/αCD28-stimulated CD8^+^T_E_) and, for reasons that remain unclear, CD8^+^T_E_ activation with PMA/ionomycin fails to elicit CCL1 protein expression with same efficacy as αCD3/αCD28 or peptide stimulation (a similar disconnect was also observed for CCL9/10 induction). Perhaps most intriguingly, our chemokine co-expression analyses revealed that the CCL1^+^ CD8^+^T_E_ subset exhibits pronounced functional diversity since this population co-produces XCL1 in addition to CCL3/4/5 and IFNγ. Beyond the visualization of chemokine co-expression patterns by FC, our results also demonstrate that induced CCL3/4/5 production by primary virus-specific CD8^+^T_E_ is co-regulated as shown by their shared compartmentalized subcellular localization and secretion in part as macromolecular complexes. While this observation is in keeping with the general capacity for complex formation by disparate chemokines [84], our analyses of chemokine-deficient mice provide additional clues for the potentially cooperative nature of T_E_ chemokine synthesis/secretion: CCL3-deficient CD8^+^T_E_, and to a lesser extent also CD4^+^T_E_, display a reduced capacity for CCL4, CCL5 and CCL9/10 production; similarly, CCL5-deficient T_E_ present with a somewhat impaired CCL3/4/9/10 response. We note, however, that we cannot rule out the possibility that these defects do not at least in part arise from the close proximity of the respective chemokine gene loci to the mutant genes in CCL3- or CCL5-deficient mice.

Arguably the most distinctive feature of the pathogen-specific T_E_ chemokine response pertains to the regulation of CCL5 production, expression and secretion; specifically, these properties comprise a delayed CCL5 production capacity of developing T_E_ populations, the constitutive CCL5 (*i.e.,* directly *ex vivo* quantifiable) expression in subcellular compartments largely distinct from cytolytic granules, and the extraordinarily fast kinetics of focused CCL5 release. In contrast to all other T_E_ activities, CCL5 expression by T cells is delayed for 3-5 days after priming as a function of regulatory control exerted by KLF13 [61–63]. Our *in vitro* experiments with LCMV-specific CD8^+^T_E_ confirm this notion (ready induction of GzmB and IFNγ but only minimal CCL5 expression within 72h of stimulation) and, to our knowledge for the first time, demonstrate these kinetics in the context of a primary CD8^+^T_E_ response *in vivo*: virtually undetectable for the first ∼3 days after LCMV challenge, constitutive CCL5 expression is found in ∼50% of specific CD8^+^T_E_ on day 5 before emerging as a property of practically all antiviral CD8^+^T_E_ by day 7-8. Thus, constitutive CCL5 expression is a distinctive hallmark for “mature” pathogen- and vaccine-specific CD8^+^T_E_ (as well as a subset of CD4^+^T_E_) that may also serve as a diagnostic readout for the better “staging” of initial T_E_ differentiation. While we did not have the opportunity to study the impact of KLF13-deficiency in our model systems, we considered another potential mechanism that may contribute to the constitutive CCL5^+^ phenotype. Human NK cells were reported to regulate constitutive CCL5 expression through the JNK/MAPK pathway [72] and in mice, NK cells are the only hematopoetic population other than T cells that presents with substantial *ex vivo* detectable CCL5 content (ref.[33] and not shown). However, as based on the undiminished constitutive CCL5 expression by NK cells or specific CD8^+^ and CD4^+^T_E_ under conditions of JNK1- or JNK2-deficiency, the JNK/MAPK pathway does not appear to contribute to the CCL5^+^ phenotype in mice.

The ready visualization of both constitutive and induced CCL5 expression by pathogen-specific CD8^+^T_E_ also may resolve seeming discrepancies pertaining to its exact subcellular distribution in human and/or murine T cell clones, blasts or primary CD8^+^T cell subsets [17, 64, 65, 69, 70]. Evaluated directly *ex vivo*, the CCL5 content of CD8^+^T_E_ is preferentially distributed across multiple vesicles discrete from GzmA- and/or GzmB-containing cytolytic granules. Yet an occasional polarization and coalescence of GzmA/B^+^ and CCL5^+^ vesicles, likely indicative of most recent T cell activation, is substantially increased following deliberate TCR stimulation, further incorporates newly synthesized CCL3/4, and thus provides a foundation for the focused release of CCL3/4/5 in part as macromolecular complexes. In fact, CCL3/4 translation by CD8^+^T_E_, just like that of IFNγ, is initiated from abundantly present mRNA templates within just 30min after TCR triggering, and is subsequently amplified by the robust induction of additional mRNA transcription. In contrast, the release of pre-stored CCL5 after TCR engagement is near-instantaneous, even precedes the full mobilization of cytolytic granules, and is primarily directed towards the IS formed between CD8^+^T_E_ and sensitized target cells. The combination of remarkably fast and focused CCL5 accumulation in a tight interaction space may temporarily create conditions associated with a spike of local CCL5 concentrations in excess of 1.0μM, *i.e.* a microenvironment that can promote conjugate stabilization (achieved, for example, already with 130nM CCL5 added to *in vitro* cultures [105]) and may contribute to receptor-independent target cell activation [80]. Interestingly, although the initial burst of CCL5 secretion is followed by additional protein production, translation is restricted to the utilization of pre-existing mRNA species since, in contrast to all other CD8^+^T_E_ chemokines, cytokines and TNFSF ligands, no further transcription is induced for at least 3h of TCR activation, and secretion of newly synthesized as opposed to pre-stored CCL5 is sensitive to inhibition by BFA; thus, CD8^+^T_E_ activation promotes two successive waves of CCL5 release characterized by their distinctive temporospatial organization of CCL5 synthesis, storage and secretion.

Again, however, it remains unclear to what extent the specific T_E_ CCL5 response and its unique characteristics may provide relevant and non-redundant contributions to the control of infectious diseases, especially in experimental or natural scenarios beyond HIV infection. For one, the historically preferred experimental usage of chemokine receptor-deficient mice complicates any interpretation pertaining to the precise role of CCL5 due to its promiscuous receptor usage (CCR1/3/5) as well as receptor-independent modes of action [106]. The use of CCL5 neutralization or CCL5-deficient mice can address these issues and although to date employed less frequently in infectious disease studies, the targeting of CCL5 collectively shows a mostly modest impairment of pathogen-specific T_E_ immunity that results, depending on experimental systems, in ameliorated immunopathology or exacerbated disease due to compromised pathogen control (reviewed ref.[106]); if any of these phenotypes are contingent on the specific lack CCL5 produced by T_E_ rather than other hematopoetic sources remains an open question.

In summary, we demonstrate that the prodigious production of chemokines, purveyors of cues essential to the coordination of complex immune responses, constitutes a circumscribed yet diverse, prominent and largely consistent component integral to the functionality of pathogen- and vaccine-specific T_E_. Further characterized by several unique aspects pertaining to the synthesis, co-expression and regulation as well as secretion of certain chemokines, the T_E_ chemokine response is readily visualized, quantified and dissected by analytical FC. As such, we propose that T cell profiling according to six distinct chemokines will considerably expand the repertoire of functional T cell assays and, importantly, may provide potentially important insights into specific T cell immunity under various experimental and naturally occurring conditions. We have pursued some of that work in series of ongoing investigations that delineate the chemokine signatures of naïve and pathogen-specific memory T cells, and under condition of prolonged antigenic persistence (manuscripts in preparation).

## MATERIALS AND METHODS

### Ethics statement

All procedures involving laboratory animals were conducted in accordance with the recommendations in the “Guide for the Care and Use of Laboratory Animals of the National Institutes of Health”, the protocols were approved by the Institutional Animal Care and Use Committees (IACUC) of the University of Colorado (permit numbers 70205604[05]1F, 70205607[05]4F and B-70210[05]1E) and Icahn School of Medicine at Mount Sinai (IACUC-2014-0170), and all efforts were made to minimize suffering of animals.

### Mice

C57BL6/J (B6), congenic B6.CD90.1 (B6.PL-*Thy1^a^*/CyJ) and B6.CD45.1 (B6.SJL-*Ptprc^a^ Pepc^b^*/BoyJ) mice; B6.CCL3^-/-^ (B6.129P2-Ccl3^tm1Unc/^J), B6.CCL5^-/-^(B6.129P2-Ccl5^tm1Hso^/J), B6.CCR5^-/-^ (B6.129P2-Ccr5^tm1Kuz^/J), B6.Jnk1^-/-^ (B6.129S1-*Mapk8^tm1Flv^*/J) and B6.Jnk2^-/-^ (B6.129S2-*Mapk9^tm1Flv^*/J) mice on a B6 background, as well as Balb/c and CCR3^-/-^ (C.129S4-Ccr3^tm1Cge^/J) mice on a Balb/c background were purchased from The Jackson Laboratory; CCR1^-/-^ (B6.129S4-Ccr^1tm1Gao^) [107] were obtained from Taconic; CCL3/CCR5-deficient mice were derived from intercrosses of B6.CCL3^-/-^ x B6.CCR5^-/-^ F1 offspring; p14 TCRtg mice on a B6.CD90.1 background were provided by Dr. M. Oldstone (CD8^+^T cells from these mice [“p14 cells”] are specific for the dominant LCMV-GP_33-41_ determinant restricted by D^b^ [35]); and B6.CCL1^-/-^ mice were a gift from Dr. S. Manes [108]. To generate p14 chimeras, naïve p14 T cells were enriched by negative selection and ∼5×10^4^ cells were transferred i.v. into sex-matched B6 recipients that were challenged 24h later with LCMV [34]; to assure reliable detection of cytolytic effector molecules [109], additional p14 chimeras were generated with lower numbers (∼10^3^) of p14 cells.

### Pathogen infections and vaccination

Lymphocytic choriomeningitis virus (LCMV) Armstrong (clone 53b) and vesicular stomatitis virus (VSV) Indiana were obtained from Dr. M. Oldstone, stocks were prepared by a single passage on BHK-21 cells, and plaque assays for determination of virus titers were performed as described [110]. Recombinant *L. monocytogenes* (LM) expressing full-length ovalbumin (rLM-OVA) [51] was grown and titered as described [111]. In brief, aliquots of ∼10^8^ mouse-passaged rLM-OVA were frozen at –80°C. To estimate titers prior to *in vivo* challenge, thawed aliquots were used to inoculate 5-10ml fresh TSB media, grown at 37°C in a shaker for 2-3h to log phase followed by determination of OD_600_ values. 8-10 week old mice were infected with a single intraperitoneal (i.p.) dose of 2×10^5^ plaque-forming units (pfu) LCMV Armstrong, 1×10^6^ pfu VSV i.v., or 3×10^2^-3×10^4^ cfu rLM-OVA i.v. as indicated; combined TLR/CD40 vaccinations were performed essentially as described [54], *i.e.* mice were immunized i.p. with 500μg ovalbumin (Sigma) or 100μg 2W1S peptide (Pi Proteomics) in combination with 50μg αCD40 (FGK4.5, BioXCell) and 50μg polyinosinic:polycytidylic acid (poly[I:C], Amersham/GE Healthcare); all vaccinations were performed by mixing each component in PBS and injection in a volume of 200μl. In some cases (***Fig.9***), mice were challenged intracerebrally (i.c.) with 1×10^3^ pfu LCMV Armstrong [27]; due to the lethal disease course in wt mice, we employed an IACUC-approved scoring matrix to measure morbidity, and terminally ill mice were euthanized and scored as deceased.

### Lymphocyte isolation, T cell purification, and stimulation cultures

Lymphocytes were obtained from spleen and blood using standard procedures [48, 112]. Splenic CD90.1^+^ p14 T_E_ from LCMV-infected p14 chimeras were positively selected using αCD90.1-PE ab and PE-specific magnetic beads (StemCell Technologies); additional purification (>99%) was achieved by FACS sorting (BDBiosciences FACS Aria). Primary cells were cultured for 0.5-5.0h in complete RPMI (RPMI1640/GIBCO, supplemented with 7% FCS, 1% L-glutamine, 1% Pen/Strep) and, where indicated, stimulated with specific peptides (1μg/ml for MHC-I- and 5μg/ml for MHC-II-restricted peptides); plate-bound αCD3 (10μg/ml) and soluble αCD28 (2μg/ml); PMA/ionomycin (5-20ng/ml and 500ng/ml, respectively); or LPS (500ng/ml, Sigma) in the presence or absence of 1μg/ml brefeldin A (BFA, Sigma). For transcriptional and/or translational blockade, cells were pre-incubated for 30min at 37°C with 5μg/ml actinomycin D (ActD, Sigma) and/or 10μg/ml cycloheximide (CHX, Sigma) prior to addition of peptide and/or BFA. *In vitro* and *in vivo* T cell proliferation was monitored by CFSE dilution as described [47, 48].

### Microarray hybridization and analysis

For microarray analyses (***Figs.1A, S2 & S3A***), p14 T_E_ were purified (>99%) from individual p14 chimeras on d8 after LCMV challenge by sequential magnetic and fluorescence activated cell sorting as detailed above; DNA-digested total RNA was extracted either directly post-sort (*ex vivo*), or after 3h stimulation with αCD3/αCD28 (see above) using a MinElute kit (Qiagen), and RNA integrity confirmed by PicoChip RNA technology (Agilent) according to manufacturer’s instructions. Amplification and labeling of mRNA (Ovation Biotin RNA Amplification and Labeling System, NuGen), hybridization to Affymetrix M430.2 arrays, and quality control were performed by the Affymetrix Core Facility of the University of Colorado Cancer Center according to standard protocols; further experimental and analytical details are provided in ref.[34], and the data can be retrieved from the GEO repository accession number GSE143632. Note that the files deposited therein also contain data for *ex vivo* p14 T_E_ previously uploaded in the context of a related study (GSE38462) as well as data on *ex vivo* and αCD3/αCD28-stimulated p14 T_M_ to be discussed in a separate manuscript in preparation (though all *ex vivo* and stimulated p14 T_E_ and T_M_ data were generated in the same set of experiments). MAS5, RMA, and GC-RMA normalization were performed and yielded essentially similar results (not shown). In addition, *ex vivo* purified and αCD3/αCD28-stimulated p14 T_E_ were analyzed by “macroarrays” (OMM022 chemokine array, SuperArray) according to protocols provided by the manufacturer and yielded results comparable to Affymetrix analyses (not shown).

### Peptides and MHC tetramers

Peptides corresponding to the indicated pathogens, ovalbumin or I-Eα epitopes were obtained from Peptidogenic, the NJH Molecular Core Facility or GenScript at purities of >95% (GP: glycoprotein; NP or N: nucleoprotein); their MHC-restriction and amino acid sequences are indicated. LCMV epitopes: GP_33-41_ (D^b^/KAVYNFATC), GP_67-77_ (D^b^/IYKGVYQFKSV), GP_92-101_ (D^b^/CSANNAHHYI), GP_118-125_ (K^b^/ISHNFCNL), GP_276-286_ (D^b^/SGVENPGGYCL), NP_396-404_ (D^b^/FQPQNGQFI), NP_166-175_ (D^b^/SLLNNQFGTM), NP_205-212_ (K^b^/YTVKYPNL), NP_118-126_ (L^d^/RPQASGVYM), GP_64-80_ (IA^b^/GPDIYKGVYQFKSVEFD), NP_309-328_ (IA^b^/SGEGWPYIACRTSIVGRAWE); VSV epitopes: N_52-59_ (K^b^/RGYVYQGL), GP_415-433_ (IA^b^/SSKAQVFEHPHIQDAASQL); rLM-OVA epitopes: OVA_257-264_ (K^b^/SIINFEKL), LLO_190-201_ (IA^b^/NEKYAQAYPNVS); and the I-Eα-derived epitope 2W1S (IA^b^/EAWGALANWAVDSA). D^b^NP_396_, D^b^GP_276_, D^b^GP_33_, L^d^NP_118_, IA^b^GP_66_ and IA^b^huCLIP_87_ complexes were obtained from the NIH tetramer core facility as APC or PE conjugates and/or biotinylated monomers; K^b^OVA_257_, IA^b^2W1S and IA^b^GP_61-80_ tetramers were prepared in the laboratory as described [47, 113, 114]. Note that the shorter sequences (GP_64-80_ and GP_66-77_) within the dominant IA^b^-restricted LCMV GP_61-80_ epitope are recognized by the same population of LCMV-specific CD4^+^T cells [115]. Tetramer staining was performed as described [36, 47].

### Antibodies, staining procedures and flow cytometry (FC)

All FC antibodies were obtained as purified, biotinylated and/or fluorochrome-conjugated reagents from RnDSystems, BDBiosciences, ebioscience, Biolegend or Invitrogen; our protocols for cell surface and intracellular FC staining, including the stringent characterization and usage of chemokine-specific monoclonal (mab) and polyclonal (pab) antibodies, are detailed elsewhere [33, 47, 112]; the utility of a new CXCL3 pab included here (RnDSystems AF5568) is demonstrated in ***Fig.S1B***. For concurrent use of two chemokine-specific goat pabs, we performed pre-conjugations with Zenon AF488 and AF647 kits according to the manufacturer’s instructions (Invitrogen). Note that the pabs αCCL1, αCCL4 and αXCL1 do not exhibit crossreactivity with any other chemokines; αCCL3 is weakly crossreactive with CX3CL1 (not expressed by any hematopoetic cells); and αCCL5 demonstrates very minor crossreactivity with CCL3 [33]. Analyses of CCL6 and CCL9/10 expression are complicated by the fact that αCCL6 and αCCL9/10 pabs exhibit significant crossreactivity with the respective non-cognate (but no other) chemokine [33]. However, since T cells fail to produce CCL6 as determined with the non-crossreactive 262016 mab (RnDSystems) [33], chemokine expression by T cells stained with the αCCL9/10 pab can be attributed exclusively to the presence of CCL9/10. Additional chemokine abs employed here include αCCL3-PE mab (clone 39624, RnDSystems) and αCCL5 mab R6G9 (mIgG_1_) generation of which has been described elsewhere [116]. For detection of murine granzymes we used GzmA clone 3G8.5 (mIgG_2b_) conjugated to FITC or PE (Santa Cruz; similar results were obtained with a rabbit anti-serum provided by Dr. M. Simon [117]) and GzmB clones GB12 (mIgG_1_) conjugated to PE or APC (Invitrogen) or GB11 (mIgG_1_) conjugated to AF647 (Biolegend). All samples were acquired on FACS Calibur or LSRII flow cytometers (BDBiosciences) and analyzed with CellQuest, DIVA (BDBiosciences) and/or FlowJo (TreeStar) software. Comprehensive functional CD8^+^T_E_ profiling (***Fig.2F***) was performed by quantification of NP_396_-specific CD8^+^T_E_ subsets expressing individual constitutive (GzmA/B, perforin) or inducible (CCL1/3/4/5/9/10, XCL1; IFNγ, IL-2, IL-3, GM-CSF; TNFα, FasL, CD40L; degranulation/killing) effector activities; primary data are found in ***Figs.2C/E***, ***4D***, ***S3B***, inducible IL-3 and FasL expression as well as degranulation were quantified as described [34], and perforin stains were performed with antibody clone S16009B (Biolegend, not shown).

### In vivo killing and CD8^+^T_E_ activation assays

*In vivo* killing assays (***Fig.S6C***) were performed as described [118]. In brief, frequencies of D^b^NP_396_^+^ CD8^+^T_E_ or p14 T_E_ in control and experimental groups of d8 LCMV-infected mice were determined prior to assay execution to assure the presence of equal specific CD8^+^T_E_ numbers; then, differentially CFSE-labeled and peptide-coated (NP_396_ or GP_33_ peptide) *vs.* uncoated CD45.1 spleen cells were transferred i.v. followed by longitudinal blood sampling (10-240min) and assessment of killing kinetics by calculating the specific loss of peptide-coated targets as a function of time after transfer; for CCL5 neutralization, mice were treated i.v. with 100μg αCCL5 (clone R6G9) or mIgG1 isotype control (clone MOPC-21, Sigma) ∼10min prior to injection of target cells. *In vivo* CD8^+^T_E_ activation assays (***Fig.8C***) were conducted according to modified protocols originally developed by Haluszczak *et al.* [119]. Here, wt and chemokine-deficient mice 8 days after LCMV infection were injected with 250μg BFA i.p. followed 30min later by i.v. injection of saline (negative control) or 100μg GP_33_ peptide; spleens were harvested 1h later, processed and immediately stained with αCD8α antibody and D^b^GP_33_ tetramers (surface) and chemokine antibodies (intracellular).

### Conjugation assays

For conjugation assays, bead-purified p14 T_E_ (CD90.1) obtained from LCMV-infected p14 chimeras (d8) were combined at a ratio of 1:1 with EL4 thymoma cells (CD90.2, magnetically depleted of a small CCR3/5 expressing subset [∼8%] and pulsed for 1h with 1μg/ml GP_33_ peptide or left uncoated followed by two washes to remove excess peptide) in pre-warmed media in a V-bottom microtiter plate, pelleted by brief centrifugation, and cultured for 20-60min. At indicated time points, cells were immediately fixed by the addition of an equal volume of 4% PFA buffer, stained for CD90.2 and cell surface CCL5, and analyzed by FC (***Fig.7B***) or confocal microscopy (***Fig.7C***).

### Chemokine & cytokine ELISAs

Quantitation of CCL3, CCL4, CCL5 and IFNγ in tissue culture supernatants or serum was performed using respective Quantikine ELISA kits and protocols provided by the manufacturer (RnDSystems) (***Figs.7D, S4A & S5B***). For evaluation of CCL3/4/5 chemokine complex formation, supernatants of NP_396_ peptide-stimulated spleen cells (d8 after LCMV, 5h stimulation, no BFA) were diluted and incubated for 2.5h at RT in plates pre-coated with 5.0μg/ml polyclonal goat IgG, αCCL3, αCCL4, or αCCL5, and absorbed supernatants were immediately analyzed for CCL3/4/5 content by standard ELISA. To determine chemokine production on a per cell basis (***Fig.7D***), FC analyses were performed in parallel to calculate the numbers of D^b^NP_396_^+^ CD8^+^T_E_ in the stimulation culture.

### Confocal microscopy

PBMC or splenocyte suspensions were prepared 8 days after LCMV infection of B6 mice. For *ex vivo* co-localization studies (***Fig.5***), CD4^-^CD19^-^NK1.1^-^ PBMCs were sorted into GzmA^-^ and GzmA^+^ populations using a MoFlow cell sorter (Beckman Coulter); for co-localization studies of CCL3/4/5 and GzmB in 5h NP_396_-peptide-stimulated splenocytes (***Fig.S5A***), cells were stained for surface and intracellular markers followed by sorting on IFNγ^+^B220^-^CD4^-^ cells using a FACSAria cell sorter (BD); and for an assessment of conjugate formation (***Fig.7C***), we employed the conjugation assay described above. Cells were resuspended in 22% BSA and spun onto glass slides (Gold Seal Micro Slides, Ultra StickTM, Cat No. 3039) for 5min at 800rpm using a cytospin (Cytospin3, Shandon) and mounted using one drop of ‘ProLong Gold reagent’ (Invitrogen) with or without DAPI and a cover slip was placed on top (No. 1 1/2, Corning). After drying overnight, slides were sealed with nail polish and stored in the dark at 4°C until acquisition. Slides were analyzed with a Leica TCS SP5 confocal laser scanning microscope equipped with an inverted Leica DMI 6000 microscope, a high performance PC TCS workstation, a 488/543/633 excitation beam splitter, a UV laser (405nm, diode 50mW), an argon laser (458/476/488/496/514nm, 100mW, attenuated to 20%), a green helium/neon laser (543nm, 1 mW) and a red helium/neon laser (633nm, 10mW) for excitation of DAPI, FITC/AF488, Cy3, PE and Cy5/APC/AF647, respectively. 2048×2048 and 1024×1024 pixel images were acquired sequentially with a 63x/N.A. 1.4 oil immersion lens at 1.9x and 5.95x zoom, respectively, resulting in respective effective pixel sizes of 63.2nm and 80.24nm. Prism spectral detectors were manually tuned to separate labels (DAPI, 415-487nm; FITC/AF488, 497-579nm; Cy3, 551-641nm; PE, 585-699nm; Cy5, 640-778nm). The pinhole size was set at 1 airy unit to give an effective optical section thickness of approximately 0.5 μm. Gray-scale images were digitized at 8 bits per channel and pseudo-colored as indicted in the figure legends using the LEICA Sp5 Software or exported as TIFF files for processing in Adobe Photoshop CS (version 8.0).

### Statistical analyses

Data handling, analysis and graphic representation was performed using Prism 4.0 or 6.0c (GraphPad Software, San Diego, CA). All data summarized in bar and line diagrams are expressed as mean ± 1 SE. Asterisks indicate statistical differences calculated by unpaired or paired two-tailed Student’s t-test and adopt the following convention: *: p<0.05, **: p<0.01 and ***: p<0.001. EC_50_ values (activation thresholds, ***Fig.2E***) were calculated by plotting the fraction of specific (IFNγ^+^) T cells as a function of peptide concentration (10^-6^-10^-11^M peptide for 5h) followed by non-linear regression analysis using appropriate data format and analysis functions in the Prism software.

## AUTHOR CONTRIBUTIONS

B.D., J.E. & D.H. designed the study; B.D., J.E., T.T.N., F.V., K.J. & V.v.d.H. conducted experiments and acquired data; R.K. provided reagents; B.D., J.E., V.v.d.H., M.V.K, A.M., R.K. & D.H. analyzed data; D.H. wrote the manuscript; and all authors contributed to manuscript editing.

## ACKNOWLEDGEMENTS

We wish to thank Dr. S. Manes for the gift of B6.CCL1^-/-^ mice; Dr. L. Lenz for rLM-OVA; Dr. T. Lane for the CCL5 antibody clone R6G9; Drs. L. Sheridan, L. Edwards and J. Humann for experimental assistance; Drs. P. Marrack and J. Kappler and the entire the “K/M lab” for critical discussion; and Dr. F. Mortari (RnD Systems) for the generous gift of the majority of chemokine antibodies utilized in this study. This work was supported by NIH grants AG026518 and AI093637, JDRF CDA 2-2007-240, a BDC P&F grant and DERC grant P30-DK057516 (DH); NIH grants U54-HL127624 and U24-CA224260 (AM); and NIH T32 training grants AI07405, AI052066 and DK007792 (BD). The funders had no role in study design, data collection and analysis, decision to publish, or preparation of the manuscript.

## SUPPLEMENTARY FIGURE LEGENDS

**Figure S1.**
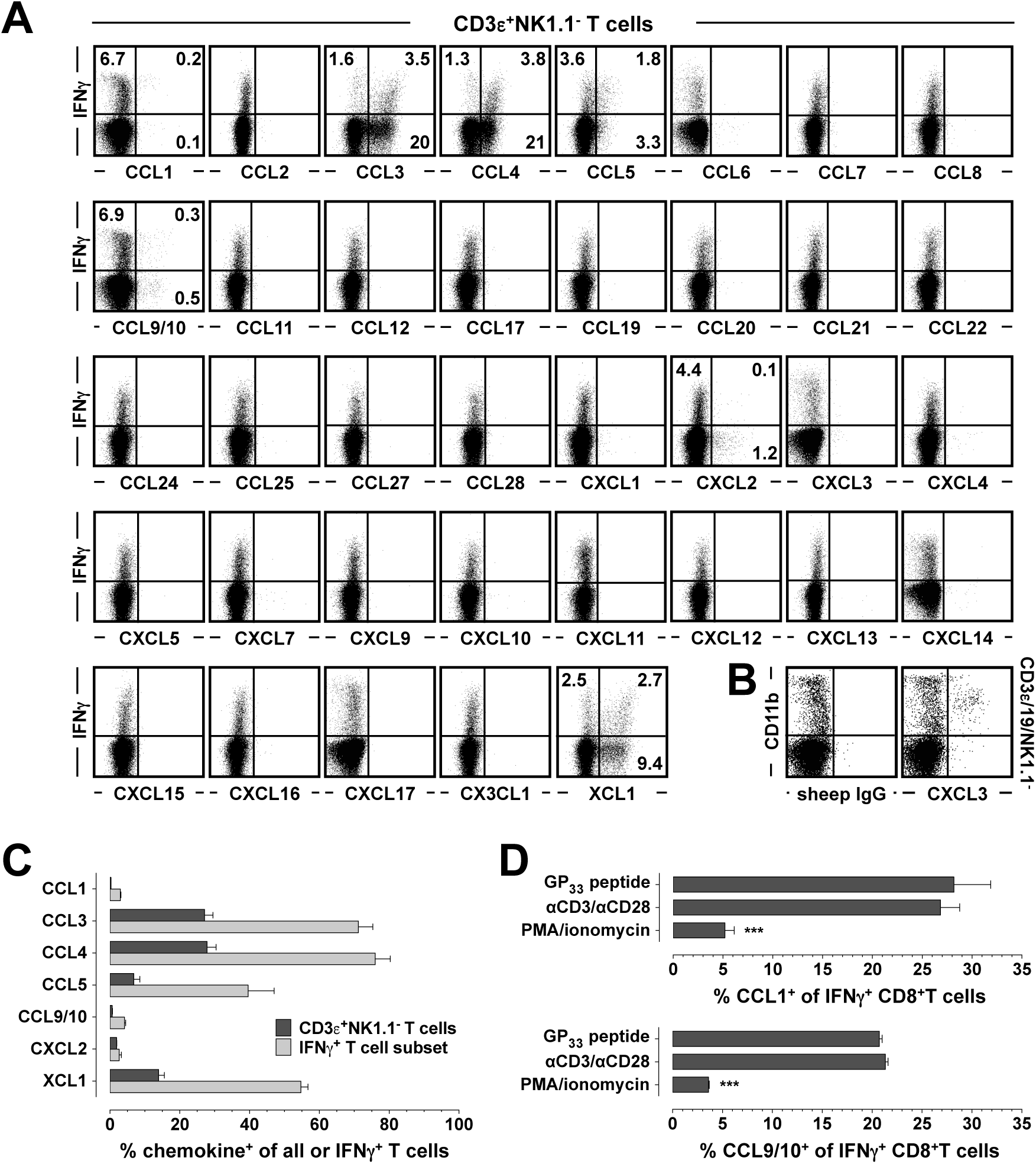
Broad survey of T cell-produced chemokines. **A.,** spleen cells from unmanipulated B6 mice were stimulated for 5h with PMA/ionomycin in the presence of BFA and stained for CD3ε, NK1.1, IFNγ and indicated chemokines as detailed in Methods (since B6 mice lack functional CXCL11, Balb/c spleen cells were used for CXCL11 expression analyses); all dot plots are gated on CD3ε^+^NK1.1^-^ cells (n=3 mice). Chemokine expression was revealed with polyclonal goat or sheep (CXCL3, CXCL17) antibodies except for CCL6 (staining performed with clone 262016 conjugated to APC) and CXCL14 (stains utilized the human CXCL14-specific clone 131120 that is crossreactive with mouse). **B.,** FC validation of the CXCL3 antibody AF5568 (polyclonal sheep, RnDSystems) used in panel A. Spleen cells were stimulated with LPS as detailed in Methods and stained for surface (CD3ε, CD11b, CD19, NK1.1) and intracellular antigens (normal sheep IgG [left] or αCXCL3 AF5568 [right]); dot plots are gated on CD3ε^-^CD19^-^NK1.1^-^ cells. **C.,** summary of T cell-produced chemokines as a fraction of all T cells (black) or the IFNγ^+^ T cell subset (n=3 mice; representative data from 3 independent experiments). **D.,** spleen cells from LCMV-immune B6 mice were stimulated with GP_33_ peptide, αCD3/αCD28 or PMA/ionomycin as described in Methods and stained for CD8α, IFNγ and CCL1 or CCL9/10. Note the substantial fraction of CCL1- and CCL9/10-producing T cells within the IFNγ compartment of GP_33_ peptide- or αCD3/αCD28-stimulated CD8^+^T cells; in contrast, PMA/ionomycin only elicited small population of CCL1^+^IFNγ^+^ and CCL9/10^+^IFNγ^+^ CD8^+^T cells (n=3 mice or triplicate samples; statistical analysis performed with one-way ANOVA).

**Figure S2.**
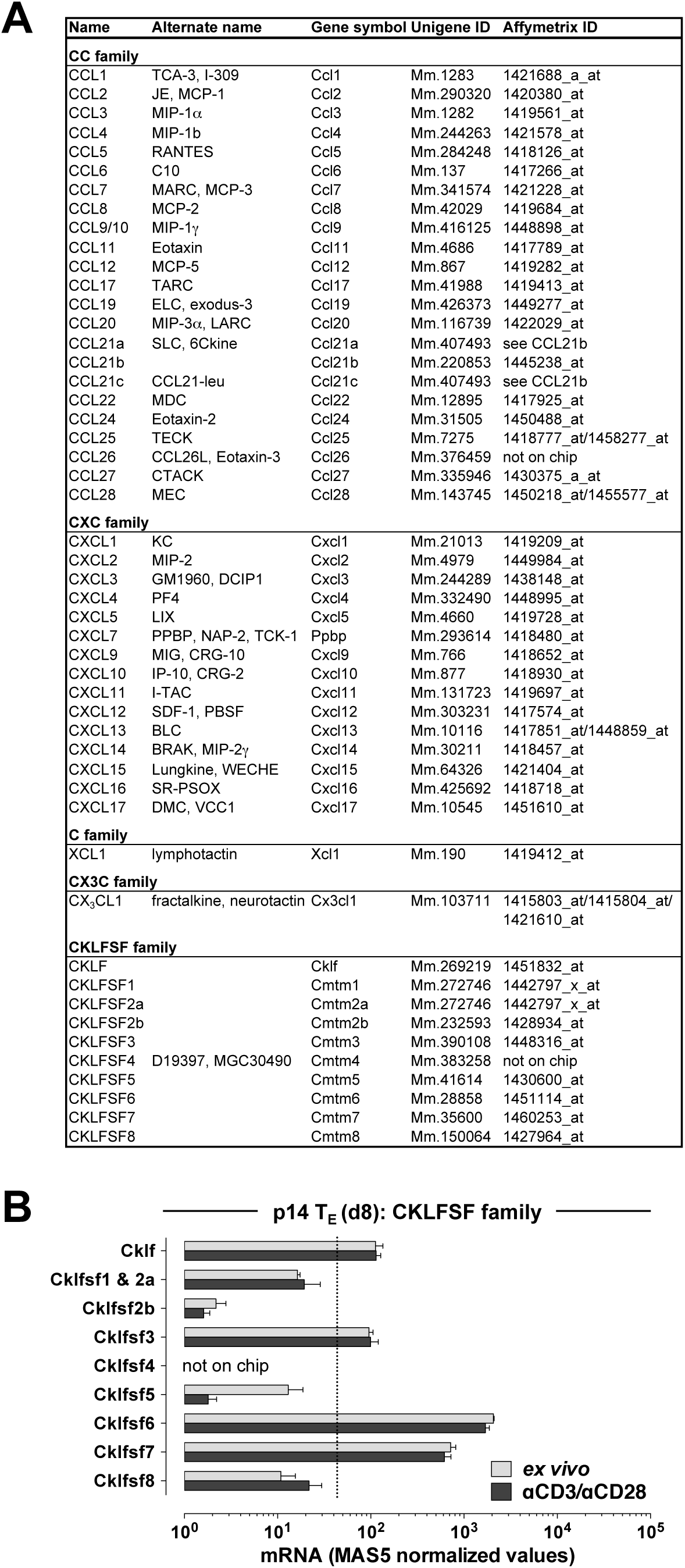
Murine chemokine nomenclature, gene and microarray IDs & CKLFSF mRNA expression by p14 T_E_. **A.,** summary of murine chemokine genes, alternative names, Unigene and Affymetrix 430 2.0 array IDs. If more than one Affymetrix ID is listed, the data presented in ***Figs.1A & S2B*** displays the average of the respective chemokine mRNA levels. **B.**, p14 CD8^+^T_E_ were purified from spleens of LCMV-challenged p14 chimeras (d8) and processed for gene array analysis as detailed in the legend to Fig.1A and Methods (n=3 individual mice). The bar diagrams display MAS5-normalized values of chemokine-like factor superfamily (CKLFSF) mRNA expression of p14 T_E_ analyzed *ex vivo* (gray bars) or after TCR stimulation by αCD3/αC28 (black bars). Statistically significant *ex vivo* or induced *Cklfsf* gene expression above the threshold of the MAS5 value of 40 (broken line) was demonstrated for *Cklf*, *Cklfsf3*, *Cklfsf6* and *Cklfsf7*. *Cklfsf4*, a gene not covered by the Affymetrix 430 2.0 array, was not detected in purified p14 CD8^+^T_E_ analyzed with SuperArray OMM022 “macroarrays” (not shown).

**Figures S3.**
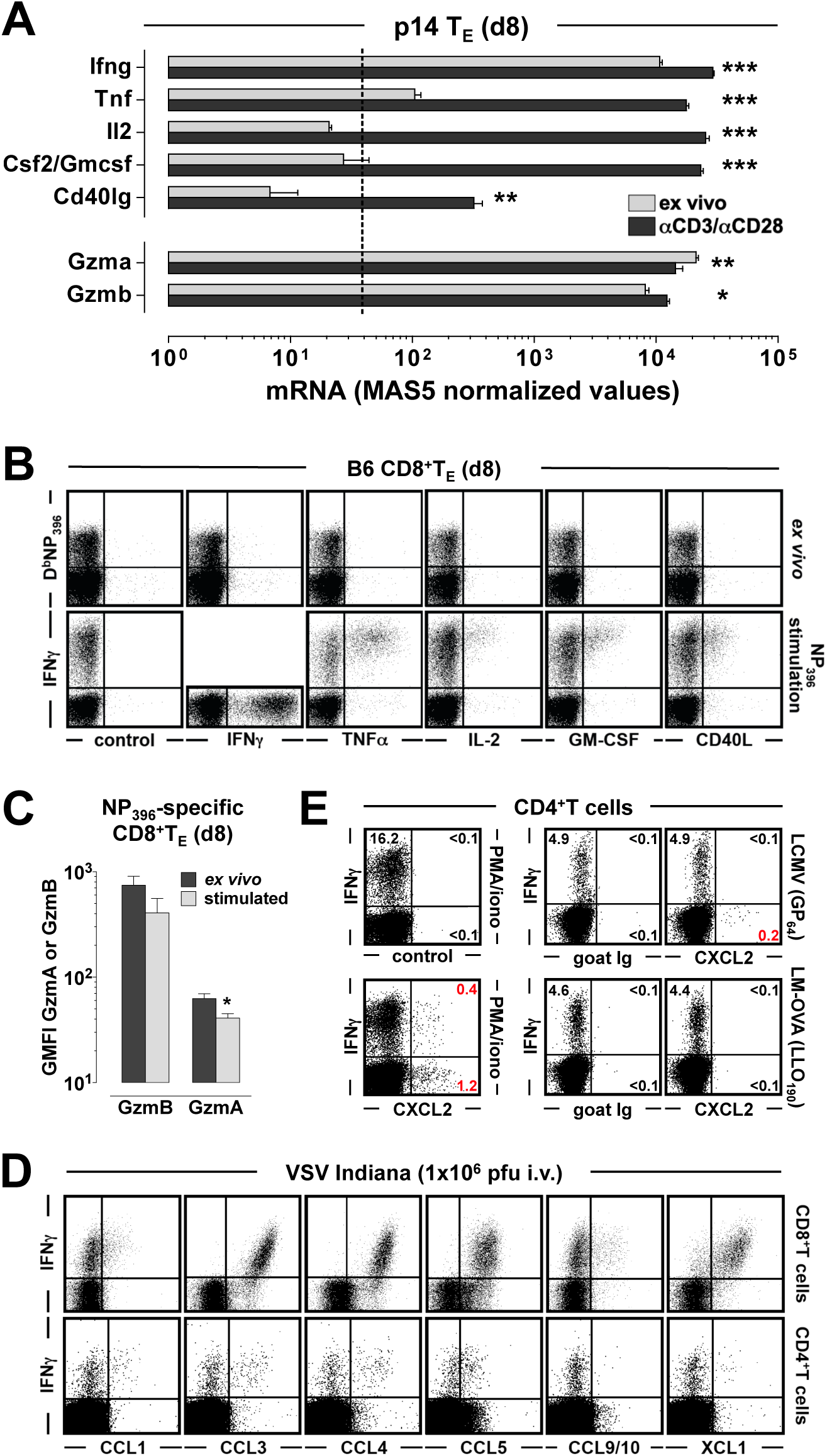
Constitutive *vs.* induced expression of CD8^+^T_E_ effector molecules & chemokine expression profiles by pathogen-specific CD8^+^ and CD4^+^T_E_. **A.**, the bar diagrams, organized as in *Figs.1A & S2B*, display the level of mRNA transcripts corresponding to major T cell-produced effector molecules expressed by LCMV-specific p14 T_E_; statistically significant differences between individual mRNA species analyzed *ex vivo* and after stimulation are indicated by asterisks (n=3 mice). **B.,** eight days after LCMV infection of B6 mice, NP_396_-specific CD8^+^T_E_ were analyzed directly *ex vivo* (top) or after 5h peptide stimulation (bottom) for the presence of major effector molecules (dot plots gated on CD8^+^T cells); note that IFNγ, TNFα, IL-2, GM-CSF and CD40L are not constitutively expressed by CD8^+^T_E_ and require a brief period of TCR stimulation to initiate protein synthesis. **C.,** in contrast, detection of constituents within the perforin/granzyme pathway does not require prior activation of specific CD8^+^T_E_ and even resulted in a slight reduction of GzmA and B expression levels after stimulation (also compare also Fig.3D; data are representative for ≥2 independent experiments performed with 3 mice/group). **D.,** B6 mice challenged with 10^6^ pfu VSV i.v. and analyzed 8 days later for induced chemokine expression by VSV N_52_-specific CD8^+^ (top) and GP_415_-specific CD4^+^T_E_ (bottom); a summary of the data is featured in Fig.2H. **E.,** left panels: spleen cells obtained from Balb/c mice were stimulated for 5h with PMA/ionomycin in the presence of BFA and subsequently stained for surface and intracellular markers; middle and right panels: B6 mice infected with LCMV Armstrong (top) or rLM-OVA (bottom) were restimulated with GP_64_ or LLO_190_ peptides and analyzed by cytokine/chemokine FC (all plots gated on CD4^+^T cells); note that LCMV-but not rLM-OVA-specific CD4^+^T_E_ contained a small subset of CXCL2-expressing “T_H_2-like” cells (∼5%).

**Figure S4.**
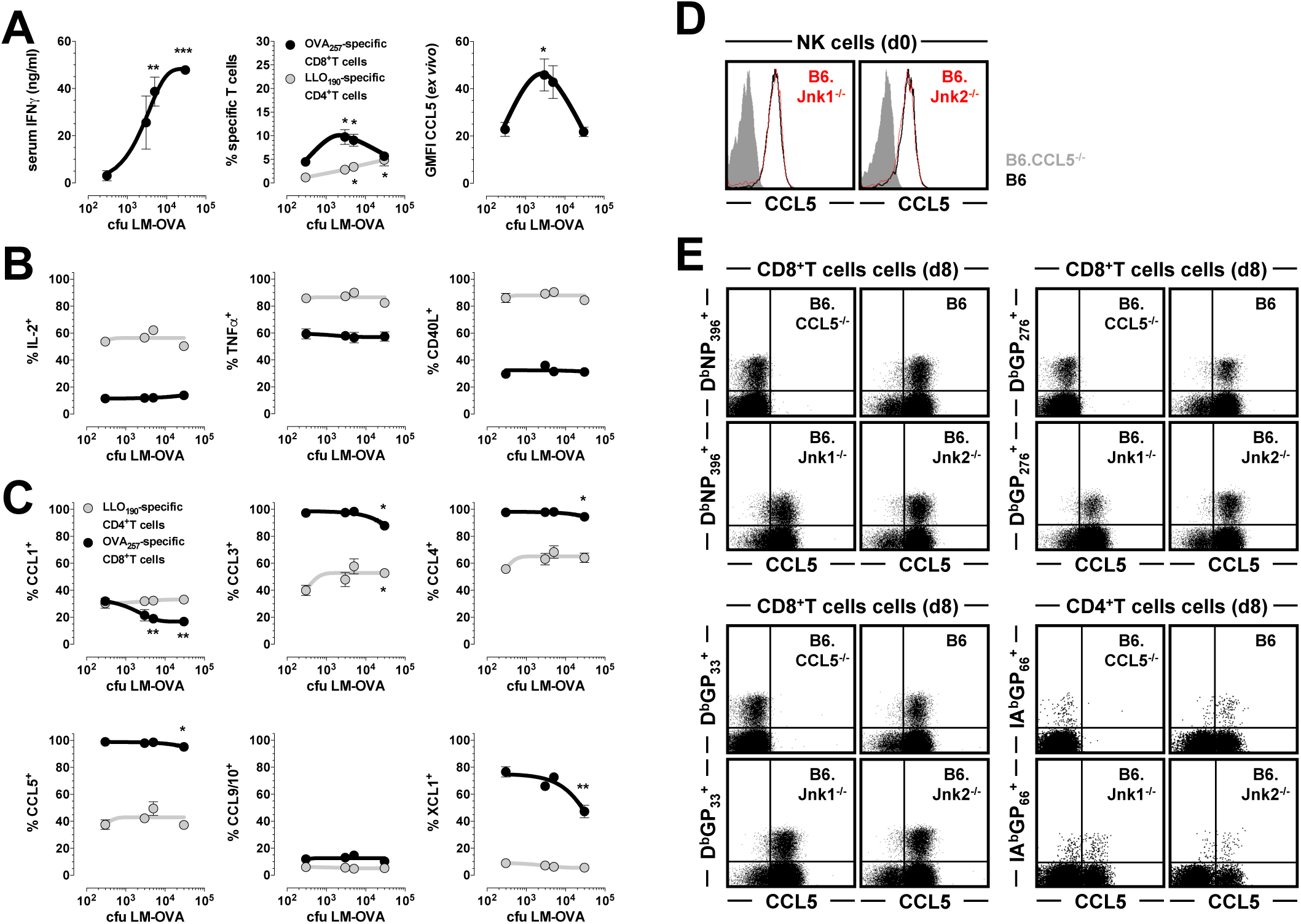
CD8^+^ and CD4^+^T_E_ chemokine production profiles as a function of rLM-OVA challenge dosage & role of JNK in the regulation of constitutive CCL5 expression by T_E_ and NK cells. **A.-C.,** B6 mice were challenged with escalating dosages of rLM-OVA (3×10^2^-3×10^4^ cfu i..v.) and analyzed eight days later by ELISA and FC (n=3). A., quantification of serum IFNγ, specific CD8^+^ and CD4^+^T_E_ frequencies in blood, and *ex vivo* detectable CCL5 content by blood-borne K^b^OVA_257_^+^CD8^+^T_E_ as a function of rLM-OVA challenge dosage. B. & C., frequencies of cytokine^+^ or chemokine^+^ OVA_257_-specific (IFNγ^+^) CD8^+^T_E_ (black) and LLO_190_-specific (IFNγ^+^) CD4^+^T_E_ (gray) in the spleen as a function of rLM-OVA challenge dosage; statistically significant differences between lowest and higher rLM-OVA dosages are indicated by asterisks (one-way ANOVA). D., *ex vivo* detectable CCL5 content of blood-borne NK cells from B6 (black), B6.CCL5^-/-^ (gray), and B6.Jnk1^-/-^ or B6.Jnk2^-/-^ (red) mice as quantified by FC. **E.,** B6, B6.CCL5^-/-^, B6.Jnk1^-/-^ and B6.Jnk2^-/-^ mice were challenged with LCMV, and CCL5 expression by specific CD8^+^T_E_ (D^b^NP_396_, D^b^GP_33_, D^b^GP_276_) and CD4^+^T_E_ (IA^b^GP_66_) in peripheral blood was visualized eight days later directly *ex vivo*.

**Figure S5.**
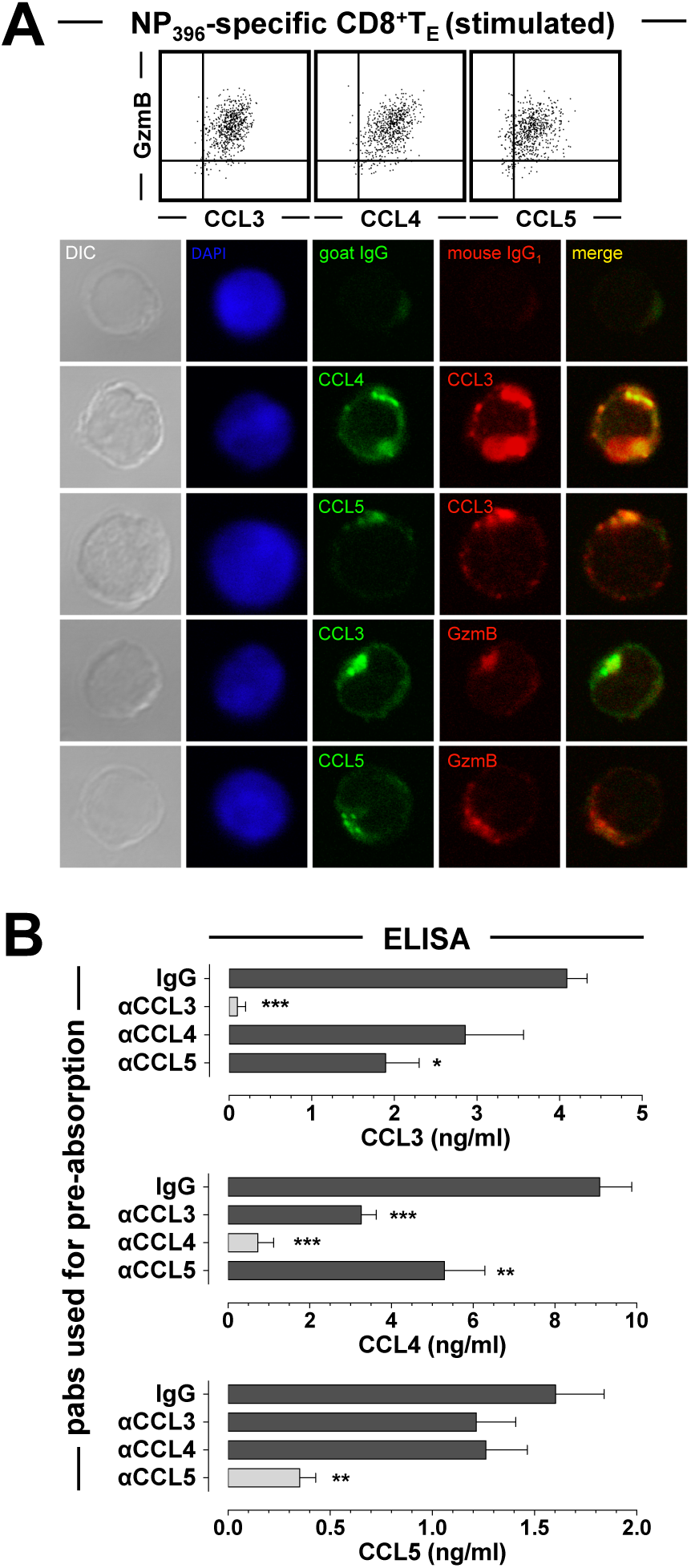
Co-expression, subcellular localization and co-secretion of CD8^+^T_E_-produced chemokines. **A.,** top: NP_396_-specific CD8^+^T_E_ (d8) stimulated for 5h with peptide in the presence of BFA to visualize GzmB and CCL3/4/5 coexpression as analyzed by FC (plots gated on NP_396-_specific [IFNγ^+^] CD8^+^T_E_). Bottom: d8 spleen cells stimulated for 5h with NP_396_ peptide in the absence of BFA to avoid interference with intracellular protein trafficking, stained for surface and intracellular markers, sorted and analyzed by confocal microscopy as detailed in Methods (negative control stains are featured in the first row); note the aggregation and co-localization of CCL3/4/5 and GzmB close to the cell membrane. **B.,** equal numbers of D^b^NP_396_^+^ CD8^+^T_E_ were peptide-stimulated for 5h (no BFA) and supernatants were pre-absorbed with indicated pabs prior to quantitation of CCL3/4/5 by ELISA (gray bars indicate absorption/detection with abs of the same specificity, statistical significance was calculated in relation to IgG pre-absorption control, and data are representative for 2 similar experiments with 3-4 mice/group).

**Figures S6.**
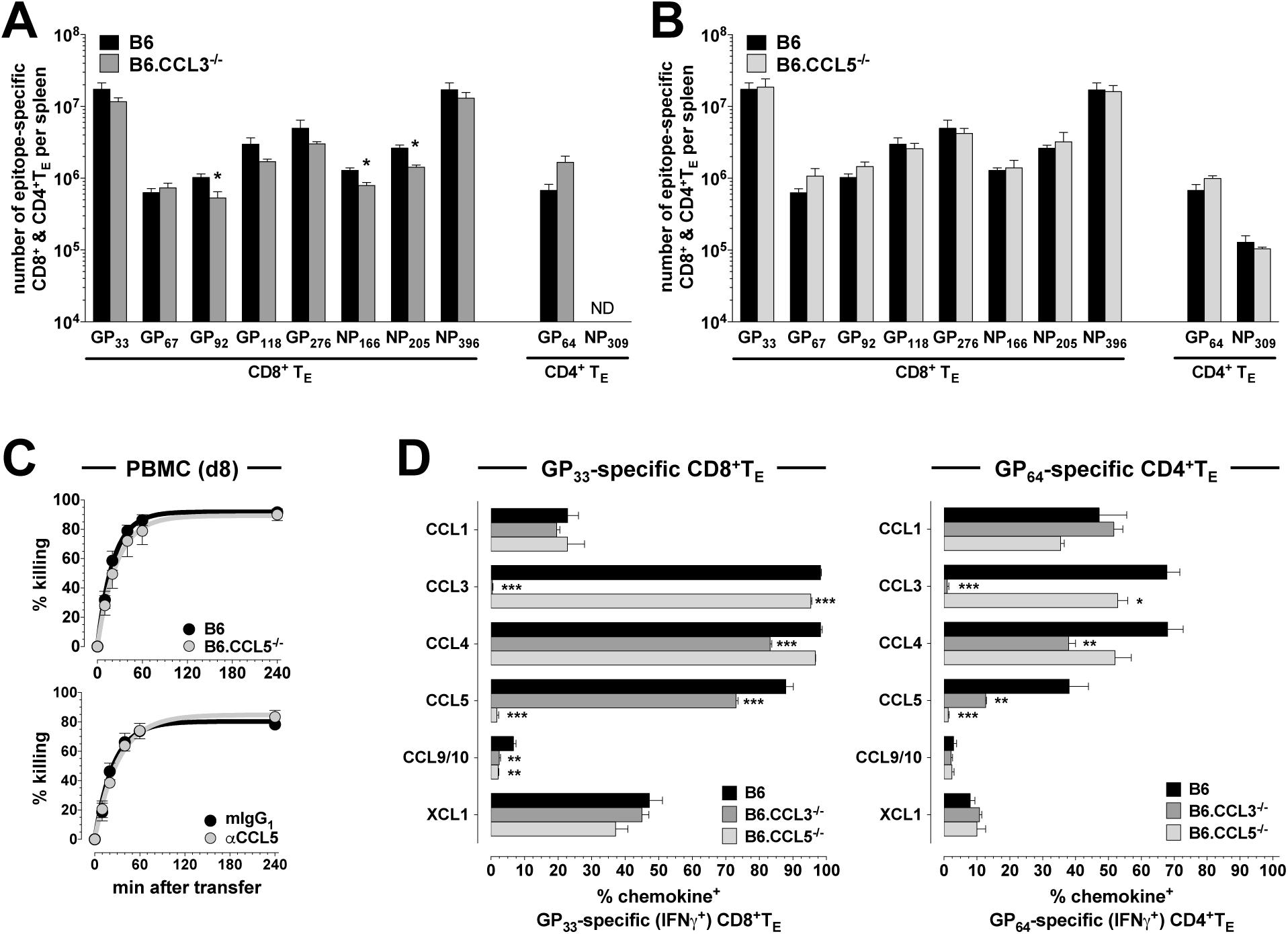
Specific T cell immunity in CCL3- and CCL5-deficient mice. **A. & B.,** B6 and B6.CCL3^-/-^ as well as B6 and B6.CCL5^-/-^ mice (n=3/group) were challenged with LCMV and numbers of epitope-specific (IFNγ^+^) CD8^+^ and CD4^+^T_E_ in the spleen were determined by FC (ND: not determined); in comparison to B6 mice, B6.CCL3^-/-^ mice exhibited a slight trend towards reduced CD8^+^ but increased CD4^+^T_E_ numbers. **C.,** top: *in vivo* killing assays with NP_396_ peptide-sensitized target cells were performed as described and referenced in Methods on d8 after LCMV infection of B6 and B6.CCL5^-/-^ mice (n=5-6 mice/group, combination of two separate experiments). Bottom: *in vivo* killing assays conducted with d8 p14 chimeras treated i.v. with 100μg αCCL5 or isotype control ∼10min prior to injection of GP_33_ peptide-coated and uncoated target cells (n=3/group). **D.,** frequencies of chemokine^+^ GP_33_-specific CD8^+^ and GP_64_-specific CD4^+^T_E_ in B6, B6.CCL3^-/-^ and B6.CCL5^-/-^ mice (n=3/group, 1/3 similar experiments; astersisks indicate significant differences between B6 and immunodeficient mice [one-way ANOVA]).

